# Life without mismatch repair

**DOI:** 10.1101/2021.04.14.437578

**Authors:** Mathijs A. Sanders, Harald Vöhringer, Victoria J. Forster, Luiza Moore, Brittany B. Campbell, Yvette Hooks, Melissa Edwards, Vanessa Bianchi, Tim H. H. Coorens, Timothy M. Butler, Henry Lee-Six, Philip S. Robinson, Christoffer Flensburg, Rebecca A. Bilardi, Ian J. Majewski, Agnes Reschke, Elizabeth Cairney, Bruce Crooks, Scott Lindhorst, Duncan Stearns, Patrick Tomboc, Ultan McDermott, Michael R. Stratton, Adam Shlien, Moritz Gerstung, Uri Tabori, Peter J. Campbell

## Abstract

Mismatch repair (MMR) is a critical defence against mutation, but we lack quantification of its activity on different DNA lesions during human life. We performed whole-genome sequencing of normal and neoplastic tissues from individuals with constitutional MMR deficiency to establish the roles of MMR components, tissue type and disease state in somatic mutation rates. Mutational signatures varied extensively across genotypes, some coupled to leading-strand replication, some to lagging-strand replication and some independent of replication, implying that the various MMR components engage different forms of DNA damage. Loss of *MSH2* or *MSH6* (MutSα), but not *MLH1* or *PMS2* (MutLα), caused 5-methylcytosine-dependent hypermutation, indicating that MutSα is the pivotal complex for repairing spontaneous deamination of methylated cytosines in humans. Neoplastic change altered the distribution of mutational signatures, particularly accelerating replication-coupled indel signatures. Each component of MMR repairs 1-10 lesions/day per normal human cell, and many thousands of additional events during neoplastic transformation.

**Highlights:**

- MMR repairs 1-10 lesions/day in every normal cell and thousands more in tumor cells
- MMR patterns and rates are shaped by genotype, tissue type and malignant transformation
- MSH2 and MSH6 are pivotal for repairing spontaneous deamination of methylated cytosine
- Replication indels and substitutions vary by leading versus lagging strand and genotype

## Introduction

Protecting the integrity of our genome is vital to the normal functioning of somatic cells and reducing their risk of transforming to cancer. The genome is under constant assault from endogenous mutagens, such as oxygen free radicals and aldehydes; from errors of DNA replication, such as those acquired in cell division or DNA repair; and from exogenous mutagens, such as ultraviolet light and chemicals in tobacco smoke. With such diverse threats to our genome, a similarly diverse set of DNA repair mechanisms have evolved, each targeting particular types of DNA damage.

The mismatch repair (MMR) pathway is a major defence against errors introduced during DNA replication by leading- and lagging-strand polymerases (Pol-*ε* and Pol-*δ* respectively). The MMR complex MutSα, comprising a heterodimer of MSH2 and MSH6, performs surveillance by sliding along histone-free DNA, engaging mismatched bases, damaged DNA or insertion/deletion loops (Kunkel and Erie, 2015). It then pairs with the MutLα complex, a heterodimer of MLH1 and PMS2, which recruits a cascade of DNA repair factors that ultimately excise a section of the lesion-containing nascent strand (Hombauer et al., 2011; Kadyrov et al., 2006). MutSα and MutLα are the major heterodimers participating in MMR, but others also contribute in eukaryotes. Perhaps most well studied is the MutSβ heterodimer of MSH2 and MSH3, which particularly targets large insertion/deletion loops and single-base-pair deletions (Johnson et al., 1996; Romanova and Crouse, 2013). It remains unclear what the relative activities of these different components are in normal human cells and during transformation.

There are also potential roles of mismatch repair in protecting genomes outside of DNA replication (Crouse, 2016; Meier et al., 2018). While there is a clear differential between template and nascent strands that guides MMR during replication, this differential is not so obvious in interphase. For a given mismatch, if MMR cannot distinguish which strand has the correct sequence, there is a risk that the repair will use the incorrect sequence as the template, fixing the mismatch as a mutation on both strands. This possibility is actually harnessed physiologically during somatic hypermutation of the immunoglobulin loci in B cells (Zubani et al., 2017), but may also introduce somatic mutations predisposing to cancer or trinucleotide repeat expansions (Peña-Diaz et al., 2012).

The detailed biochemistry of MMR pathways has been pieced together from *in vitro* experimental studies and animal models (Johnson et al., 1996; Lujan et al., 2014; Meier et al., 2018; Romanova and Crouse, 2013; Serero et al., 2014; St Charles et al., 2015). What role MMR plays in humans *in vivo* has largely been inferred from extracting mutational signatures from cancers with deficiency of one component (Alexandrov et al., 2020; Campbell et al., 2017; Chung et al., 2020; Meier et al., 2018). This has identified several signatures associated with MMR deficiency, and has also demonstrated the role of MMR in protecting strategically important regions of the genome (Frigola et al., 2017; Supek and Lehner, 2015). However, these studies have focused on somatic mutations in established cancers, meaning it can be difficult to disentangle consequences of increased cell division rates, the emergence of additional mutational signatures during transformation and other endogenous mutational processes active before MMR deficiency was acquired (Alexandrov et al., 2020; Haradhvala et al., 2018; Meier et al., 2018).

Common to both experimental studies of MMR and analyses of human cancer is the utility of studying mutational signatures in knock-outs of individual pathway components. MMR deficiency in human cancers is mostly either sporadic (both alleles somatically mutated or silenced) or inherited as an autosomal dominant condition (Lynch syndrome), where one allele is inactivated in the germline and the other allele lost somatically in the tumor. Rarely, however, individuals are born with both alleles of an MMR gene inactivated in the germline, known as constitutional mismatch repair deficiency (CMMRD). These individuals have a substantially increased risk of early-onset cancers, especially cancers of the brain, gastrointestinal system and blood (Shlien et al., 2015; Tabori et al., 2017). Here, we harnessed recently developed protocols to identify mutations in normal somatic cells (Brunner et al., 2019; Ellis et al., 2020; Lee-Six et al., 2019) to evaluate individuals with CMMRD. Studying lifelong mutation accumulation in normal cells, in neoplastic lesions, in different organ systems and in different genotypes provides direct quantification of the importance of MMR in day-to-day life and the types of DNA lesion repaired by individual pathway components in humans.

## Results

### MMR Pathways Repair 1-10 Lesions/Day in Every Intestinal Epithelial Cell

Ten CMMRD donors, aged between 3 and 32 years, were studied, comprising 2 with *MSH2*, 4 with *MSH6*, 2 with *MLH1* and 2 with *PMS2* biallelic germline deficiency (**Table S1**). Colonic (n=7), small bowel (n=6) and brain (n=3) samples were collected during endoscopy, surgical resection or autopsy. Matched normal and neoplastic intestinal epithelium samples (adenomas or adenocarcinomas) were available for 4 individuals.

Intestinal crypts contain 5-15 stem cells at their base, with single stem cells frequently sweeping to fixation through genetic drift (Lopez-Garcia et al., 2010). As a result, all epithelial cells in individual crypts derive from a single, recent common ancestor, meaning that somatic mutation burdens, rates and signature composition can be accurately estimated by sequencing single crypts (Lee-Six et al., 2019). We therefore isolated 80 normal intestinal crypts, 30 neoplastic intestinal crypts and 25 samples from other tissues by laser capture microdissection, followed by whole genome sequencing (WGS) at a median of 30-fold coverage to investigate somatically acquired mutations. We called single base substitutions (SBS), small insertions and deletions (indels), structural variants (SV), telomere length and copy number variations (CNV) using well-established bioinformatic algorithms (Campbell et al., 2008; Jones et al., 2016; Raine et al., 2015) (**Table S2**). These data were compared to previously published analyses of somatic mutations in normal intestinal epithelium from mismatch repair-proficient individuals (Lee-Six et al., 2019) (**Figure S1**).

The burdens of single base substitutions were considerably increased in the normal intestinal epithelium of CMMRD donors, across all genotypes, compared to healthy individuals (**Figure 1a**). Even with the young age of our cohort, individual normal intestinal epithelial cells frequently had tens of thousands of mutations. The SBS mutation rate varied extensively between individuals, ranging between 250 and 1450 SBS/year (**Figure 1b**), which is 5 to 30-fold higher than the 49 SBS/year rate observed for normal colonic epithelium. However, within an individual, SBS mutation burdens exhibited limited variation, suggesting that genotype and environmental modifiers are stronger influences on mutation rates than cell-to-cell variation.

**Figure 1.**
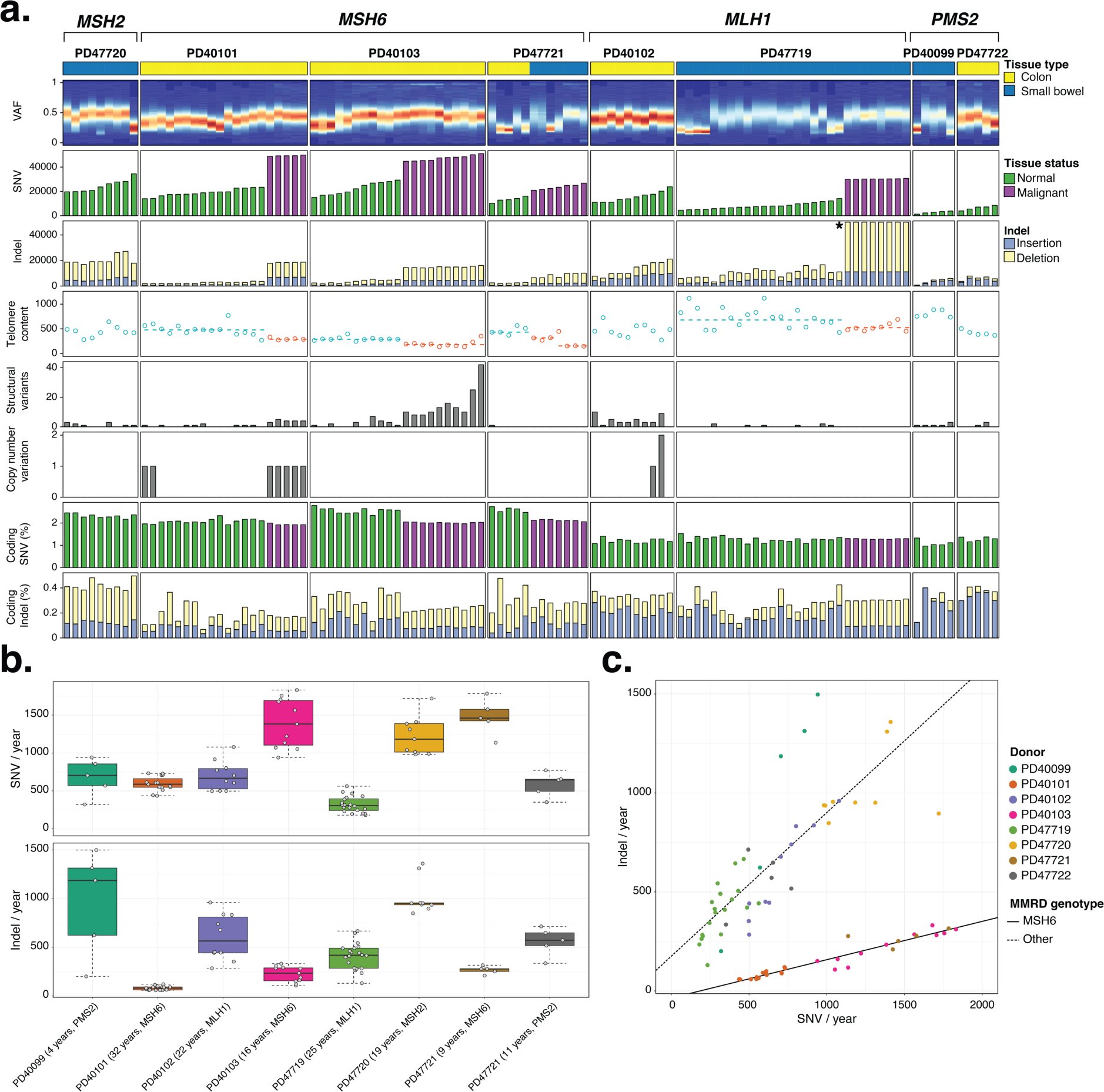
Mutation Burden and Acquisition rates in Normal and Neoplastic CMMRD Intestinal Crypts. **(a)** Each column represents an individual sample. From top to bottom: tissue type indicator, VAF distribution heatmap, SNV burden coloured by disease state, indel burden separated on insertions and deletions, telomere content, SV burden, CNV burden, percentage of coding SNVs coloured by disease state, percentage of coding indels separated on insertions and deletions. Dashed horizontal lines indicate median telomere content for matched normal and neoplastic intestinal tissue samples. **(b)** Annual SNV and indel rate for normal intestinal epithelium samples of each CMMRD donor (samples from neoplastic tissues not shown). Boxplots mark median, inter-quartile range (IQR) and whiskers extend to 1.5 IQR from last quartile. Donor ID, age and MMRD genotype are displayed below. **(c)** Relationship between annual SNV and indel rates. Individual samples are colour-coded based on donor origin. Solid line regresses the relationship for MSH6-deficient samples only. Dashed line regresses the relationship for all other MMRD genotypes. **\***, y-axis is limited to 50,000 indels to accommodate visualisation of the indel burden for all samples.

Small indel mutation burdens were similarly increased in the normal intestinal epithelium of all CMMRD donors, with an even greater magnitude of increase than for base substitutions. Rates ranged between 60 and 1200 indels/cell/year (**Figure 1b**), compared to 1 indel/cell/year for normal intestinal epithelium without MMR deficiency. As seen for base substitutions, between-individual variation was more pronounced in non-transformed epithelial cells than within-individual variation, again attributable to genotype. Comparing the number of base substitutions to indels, we found that *MSH2*, *MLH1* and *PMS2*-deficient cases all exhibited a ratio of approximately 1:1 – namely, one indel is acquired for every base substitution acquired in these normal cells (compared to 1 indel for every 49 substitutions in MMR-proficient normal epithelium; **Figure 1c**). For *MSH6* deficiency, the relative mutation rate was similarly fixed between the two types of lesions, but with a ratio of 1 indel for every 4 substitutions. The lower rate of indels relative to substitutions for *MSH6* deficiency indicates that the MutSβ heterodimer (MSH2 and MSH3) repairs ∼3x as many indels as MutSα (MSH2 and MSH6) in normal human intestinal cells.

Comparing the ratio of deletions to insertions, we found that *MSH2* and *MSH6*-deficient intestinal epithelium has a predilection for deletions, especially in polyA/T tracts, while in *MLH1*-deficient cases this ratio is at near-equilibrium and for *PMS2*-deficient cases insertions at polyA/T tracts are the predominant form of indel (**Figure S2a-b**). Upon neoplastic change the burden of deletions considerably outnumbers the insertion burden irrespective of MMRD genotype indicating that increased microsatellite instability (MSI) is predominantly defined by deletions, often at poly-A/T tracts (**Figure S2c**).

Comparing synchronous normal and neoplastic intestinal epithelium from the same individual showed that SBS burdens were 2 to 4-fold higher in neoplastic than normal intestinal epithelium, while indel burdens increased 4 to 10-fold (**Figure 1a**). As expected, telomere lengths of the intestinal crypts of neoplastic biopsies were significantly reduced compared to their normal counterpart (median normal and neoplastic telomere content, Mann-Whitney U test; PD40101, 478 vs. 287, *P*=0.0025; PD40103, 288 vs 181, *P*=0.0008; PD47719, 727 vs 514, *P*=0.008; PD47721, 432 vs 153, *P*=0.016) – this suggests that the increased cell division associated with malignant change drives both telomere shortening and increases in replication-associated somatic mutation.

Overall, then, in normal intestinal epithelial cells from patients lacking MMR, the combined rates of base substitutions and indels ranged from 500 to 3000 mutations per year per clone – this suggests that the mismatch repair pathway typically repairs an average of 1-10 events per day in every normal epithelial cell throughout the intestine.

### MMR Repairs Replication-dependent and Independent DNA Damage

The mismatch repair pathway has variable efficiency for correcting different types of mismatch, being particularly tuned to events not already targeted by the proof-reading capacity of Pol-δ and Pol-ε polymerases (Lujan et al., 2014). Thus, we might expect MMR deficiency to be associated with distinct mutational signatures, with mutation rates varying by specific base changes, indels, local sequence context and replication strand. To assess this, we extracted mutational signatures jointly for all microdissected samples, including both healthy and neoplastic, using the TensorSignatures algorithm (Vöhringer et al., 2020). TensorSignatures extracts mutational signatures from a broad set of local and global sequence information enabling the coupling of signatures of substitutions with indels as well as replication and transcriptional strand biases.

In total, 6 mutational signatures were extracted from the sequencing data of all microdissected samples, which we designate as signatures MMR-1 to MMR-6 (**Figure 2**, **Table S3**). Although these signatures partially match signatures previously extracted from cancer genomes and experimental models with MMR deficiency (Alexandrov et al., 2020; Meier et al., 2018; Serero et al., 2014; St Charles et al., 2015), the use of normal and transformed clones from cases with congenital deficiency enables clearer delineation of patterns, and isolation of individual components of previously inseparable signatures.

**Figure 2.**
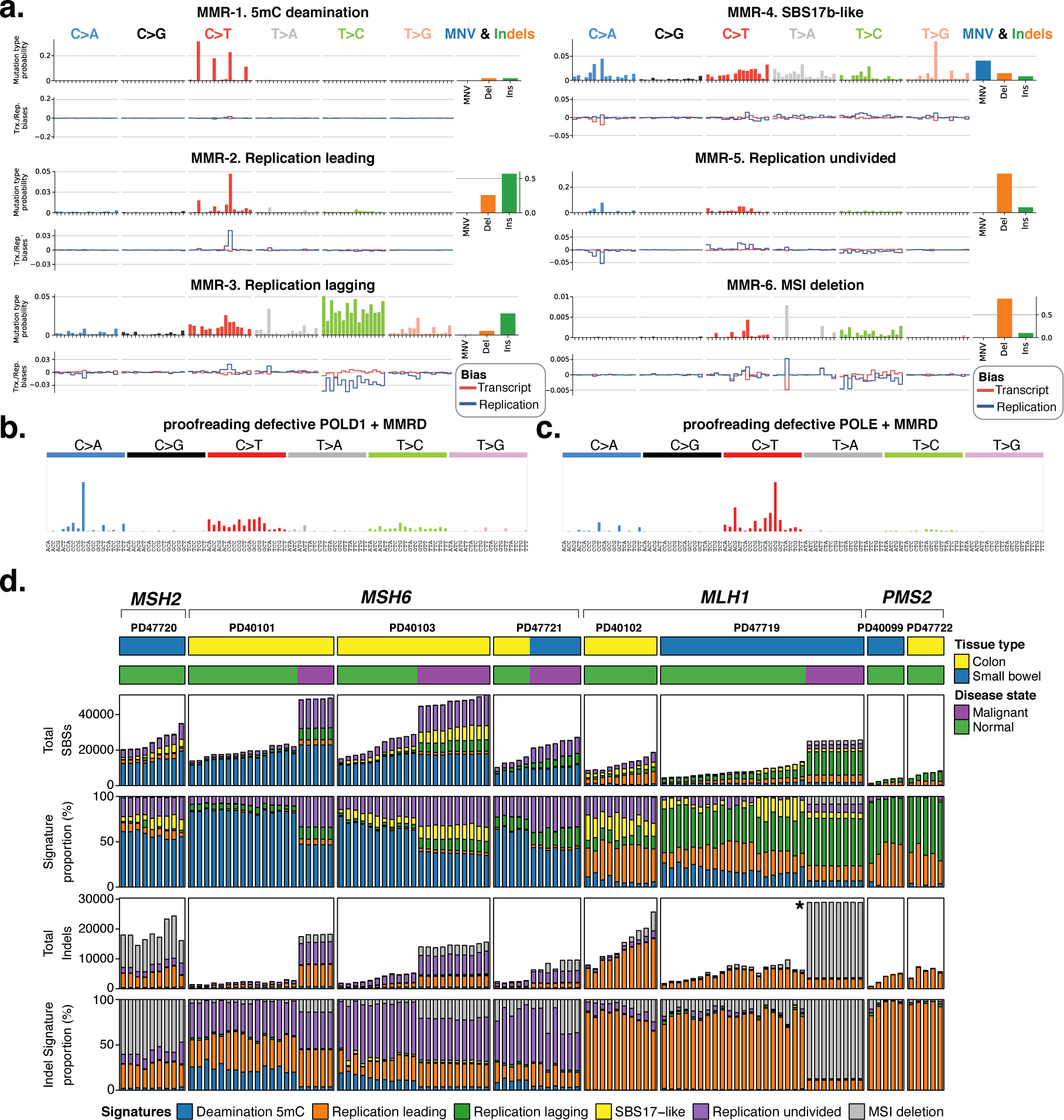
Mutational Signature Analysis Reveals DNA Damage Repair Specificity with Respect to MMRD Genotype and Disease State. **(a)** The 6 mutational signatures extracted by TensorSignatures from all CMMRD intestinal crypts and other tissues samples. Mutational catalogues for intestinal crypts were extracted from variant phylogenies (Figures 2-3) to prevent duplication of variants presents in multiple related samples. For each signature, the left top panel shows probability for the 96 possible single base substitutions classified by trinucleotide contexts grouped into the 6 possible pyrimidine-based mutations. The right top panel shows mutation type probability for MNVs, insertions and deletions. Total contribution of all possible SNVs and other mutations sums to 1. The bottom panel shows transcriptional (red line) and replication (blue line) strand biases for each possible SBS type and context. Positive values indicate coding and leading strand biases, respectively. Negative values indicate template and lagging strand biases, respectively. **(b)** Typical SBS mutational profile for cancers from TCGA marked by both proofreading defective POLD1 and concomitant loss of MMR. **(c)** Typical SBS mutational profile for cancers from TCGA marked by both proofreading defective POLE and concomitant loss of MMR. **(d)** Each column indicates an individual sample. From top to bottom: tissue type indicator, mutational signature attribution for SBSs, proportional mutational signature attribution for SBSs, mutational signature attribution for indels, proportional mutational signature attribution for indels. **\*:** y-axis is limited to 30,000 to accommodate the visualisation of mutational signature attribution for all samples.

Signature MMR-1 is characterised by C>T substitutions at <u>C</u>G dinucleotides, likely due to deamination of 5-methylcytosine (5mC). It closely matches signature SBS1, extracted from cancer genomes (Alexandrov et al., 2020), which has previously been linked with MMR deficiency (Meier et al., 2018; Supek and Lehner, 2015). MMR-1 has minimal contribution from indels and lacks transcription and replication strand biases. It was broadly present across all samples but was especially prominent in patients with defects in MutSα, namely *MSH2* or *MSH6* deficiency (**Figure 2d**, **Table S4**). Deamination of 5mC to thymine occurs as a spontaneous chemical reaction in DNA (Shen et al., 1994) throughout the cell cycle. Such events occurring in interphase require repair <u>before</u> DNA replication, otherwise the new thymine becomes fixed as a C>T mutation in the genome. The predominance of this signature in patients with MutSα deficiency suggests that a major role of the MSH2/MSH6 complex is to identify G:dT mismatches for repair in interphase, independent of its activities during replication. The paucity of this signature in the normal and neoplastic tissues from *MLH1* and *PMS2*-deficient patients implies that the repair occurs through pathways independent of MutLα, likely through nucleotide or base excision repair.

Signature MMR-2 is defined by both base substitutions and indels, predominantly A/T insertions at polyA/T tracts, with the base substitutions being specifically biased towards leading-strand replication (using the pyrimidine of each base-pair as the reference base). Signature MMR-3 is its counterpart, with substitutions being more frequent on the lagging strand of replication, comprising largely C>T and T>C transitions. In contrast to MMR-1, signatures MMR-2 and MMR-3 were most enriched in those patients with defects in MutLα (*MLH1* or *PMS2*). The distribution of base substitutions in MMR-3 closely matches signature SBS26 extracted from cancer genomes, whereas MMR-2 has only partial similarity to SBS15 (Alexandrov et al., 2020).

Signature MMR-4 strongly resembles a combination of SBS17a and SBS17b as seen in cancer genomes of the GI tract, with high rates of T>C in a C<u>T</u>T context. This signature has been associated with 5-fluorouracil treatment and misincorporation of 8-hydroxy-dGTP opposite adenines (Christensen et al., 2019; Pich et al., 2019; Supek and Lehner, 2017). Signature MMR-5 has partial similarity to SBS20 distinguished by C>A and C>T base substitutions and deletions. Interestingly in this signature, C>T events are biased towards leading-strand replication and C>A events towards lagging-strand replication. Finally, signature MMR-6 is nearly completely defined by A/T-base deletions at polyA/T tracts. It is more pronounced in *MSH2* deficiency than *MSH6* deficiency, suggesting it may capture the repair activities of the MutSβ heterodimer (MSH2 and MSH3).

The observed replication strand biases of MMR-2, MMR-3 and MMR-5 could in principle be of opposite strandedness when the complementary purine bases are considered as the reference, such as G>A on the leading strand instead of C>T on the lagging strand. Further insight can be derived from cases with combined MMRD and proofreading-defective replication polymerases from The Cancer Genome Atlas (TCGA). Given that Pol-δ and Pol-ε are the primary polymerases for lagging and leading-strand replication respectively, any overlap in mutational signatures observed in these cancers and those extracted from the CMMRD microdissections would provide evidence of their replicative origin. Cancers with proofreading-defective POLD1 combined with MMRD have distinctive mutation profiles with a predilection for C>A mutations at C<u>C</u>N as well as C>T and T>C transition mutations (**Figure 2b**). These mutational patterns resemble a combination of signatures MMR-3 and the C>A component of MMR-5 with lagging-strand bias. Cancers with proofreading-defective POLE combined with MMRD have mutation profiles with strong predilection for C>T mutations predominantly at G<u>C</u>G trinucleotides, but also G<u>C</u>N (**Figure 2c**). MMR-2 and the C>T component of MMR-5 with leading-strand bias recapitulate these mutational patterns. Taken together, then, these data suggest that signature MMR-2 is indeed tied to leading-strand replication; MMR-3 to lagging-strand replication; and MMR-5 has mixed contributions from leading and lagging-strand replication.

### Mutational Processes Vary with Genotype and Cellular Transformation

We reconstructed phylogenetic trees showing clonal relationships among normal and neoplastic intestinal epithelium crypts (n=110). In the normal epithelium of MutSα-deficient CMMRD cases (3 *MSH6*; 1 *MSH2*), the predominant mutational process was that of methylated cytosine deamination, causing C>T mutations at <u>C</u>G dinucleotides (signature MMR-1; **Figure 2a** and **3**). For example, for case PD40101, the median contribution of this signature exceeded 80%, resulting in an average 16,000 methylation-deamination mutations per cell over 32 years of life. A corollary to the predominance of the deamination signature, MMR-1, in these normal epithelial cells is that the replication-dependent MMR signatures contributed considerably fewer mutations. In contrast, in synchronously collected intestinal polyps and colorectal cancers from these same patients, the replication-based signatures, MMR-2, MMR-3 and MMR-5, became much more evident, accounting for the overwhelming majority of excess mutations seen with cellular transformation. Thus, repair of methylated cytosine deamination is numerically the most important activity of MutSα in normal intestinal epithelium, whereas repair of replication-mediated errors becomes a much more significant activity with neoplastic transformation.

**Figure 3.**
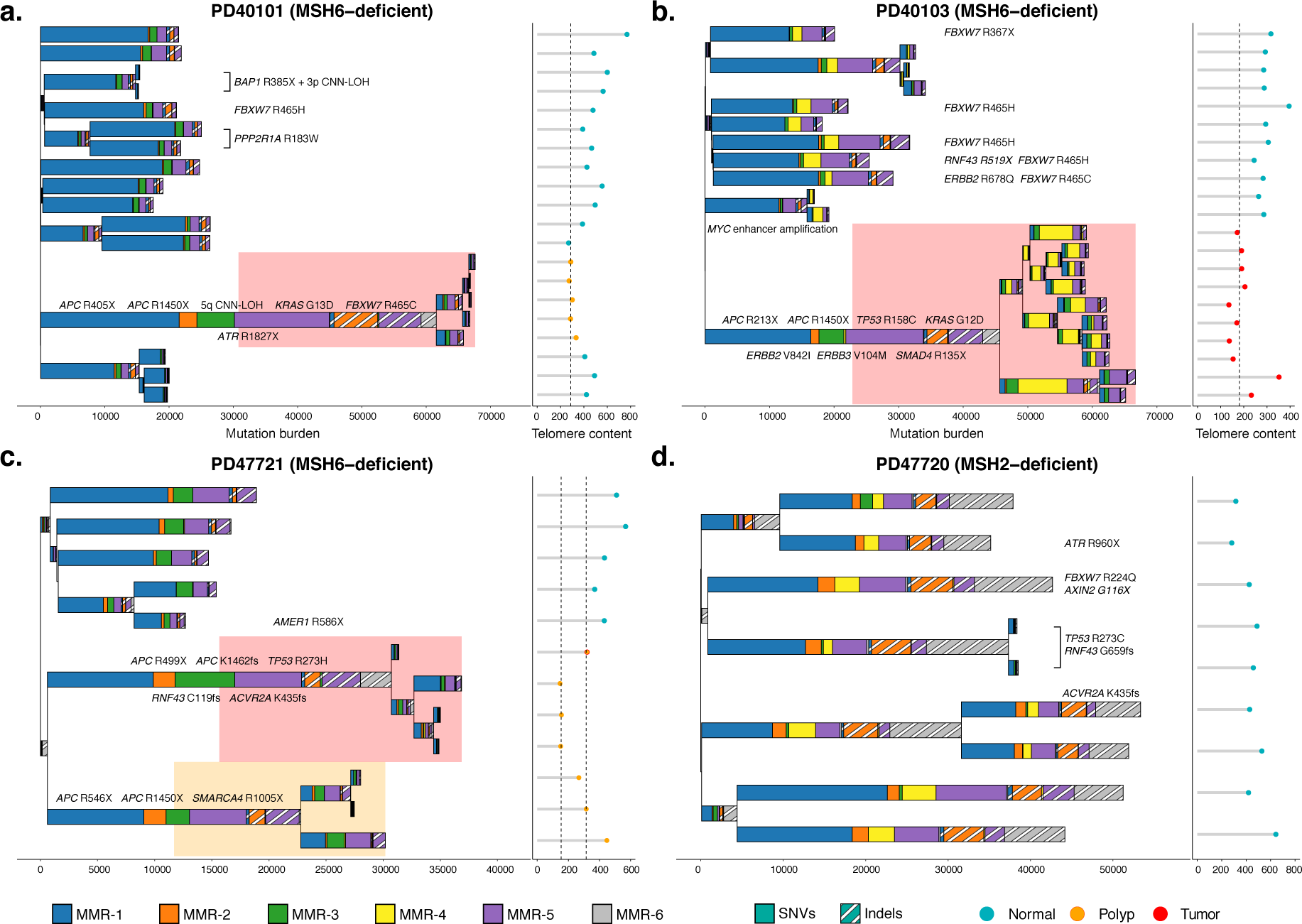
Phylogenetic Mutational Signature Attribution reveals Differential DNA Damage Repair Deficiency in Healthy and Neoplastic MutSα-deficient Intestinal Crypts. (**a-d**) Intestinal crypt phylogenies were inferred from all mutations (SNVs and indels). Branch length is proportional to the number of mutations detected. Branching indicates a historical crypt fission event. Stacked bar plots represent the mutational signature composition contributing to individual branches and colour-coded according to activity. Branches defined by <100 mutations were omitted from signature extraction. Stacked bar plot constituents: solid pattern – SNVs, stripped pattern – indels. Right panel: telomere content for individual crypts coloured by disease state. Dashed line indicates the median telomere content for neoplastic intestinal crypts. Coloured rectangular areas highlight branches formed by intestinal crypts microdissected from neoplastic tissues. Detected driver mutations are listed next to or along branches to indicate their presence among related intestinal crypts. **(a)** PD40101 (MSH6-deficient), phylogeny comprising normal (n = 12) and matched neoplastic intestinal crypts (n = 5). **(b)** PD40103 (MSH6-deficient), phylogeny comprising normal (n = 11) and matched neoplastic intestinal crypts (n = 10). **(c)** PD47721 (MSH6-deficient), phylogeny comprising normal (n = 5) and matched neoplastic intestinal crypts (n = 4, n = 3). Coloured area indicates the branches common to the 2 individual microdissected matched polyps. **(d)** PD47720 (MSH2-deficient), phylogeny comprising only normal intestinal crypts (n = 9).

Loss of MutLα showed a different distribution of mutational signatures in normal intestinal crypts compared to MutSα-deficiency (**Figure 4**). MMR-2, marked by A/T-base insertions and leading-strand C>T mutations, was common to all MutLα-deficient intestinal crypts sequenced. MMR-3, with C>T and T>C transitions biased towards the lagging-strand, was similarly present in all intestinal crypts, but was more pronounced in *PMS2*-deficient than *MLH1*-deficient cases. This suggests that even among cases with deficiencies of components from the same MMR complex there are differences in the relative activity of mutational processes. Neoplastic change also impacted the composition of operative mutational processes, as seen in PD47719 (**Figure 4a**). Here, MSI was considerably increased in the transformed clone, as evidenced by the massive, sustained contribution of MMR-6 to the excess mutations seen on this branch of the phylogeny, including the most recent branches. The consequence of this was that, while insertions predominated over deletions in the normal epithelium of MutLα-deficient patients including PD47719, deletions considerably outnumbered insertions in this transformed clone.

**Figure 4.**
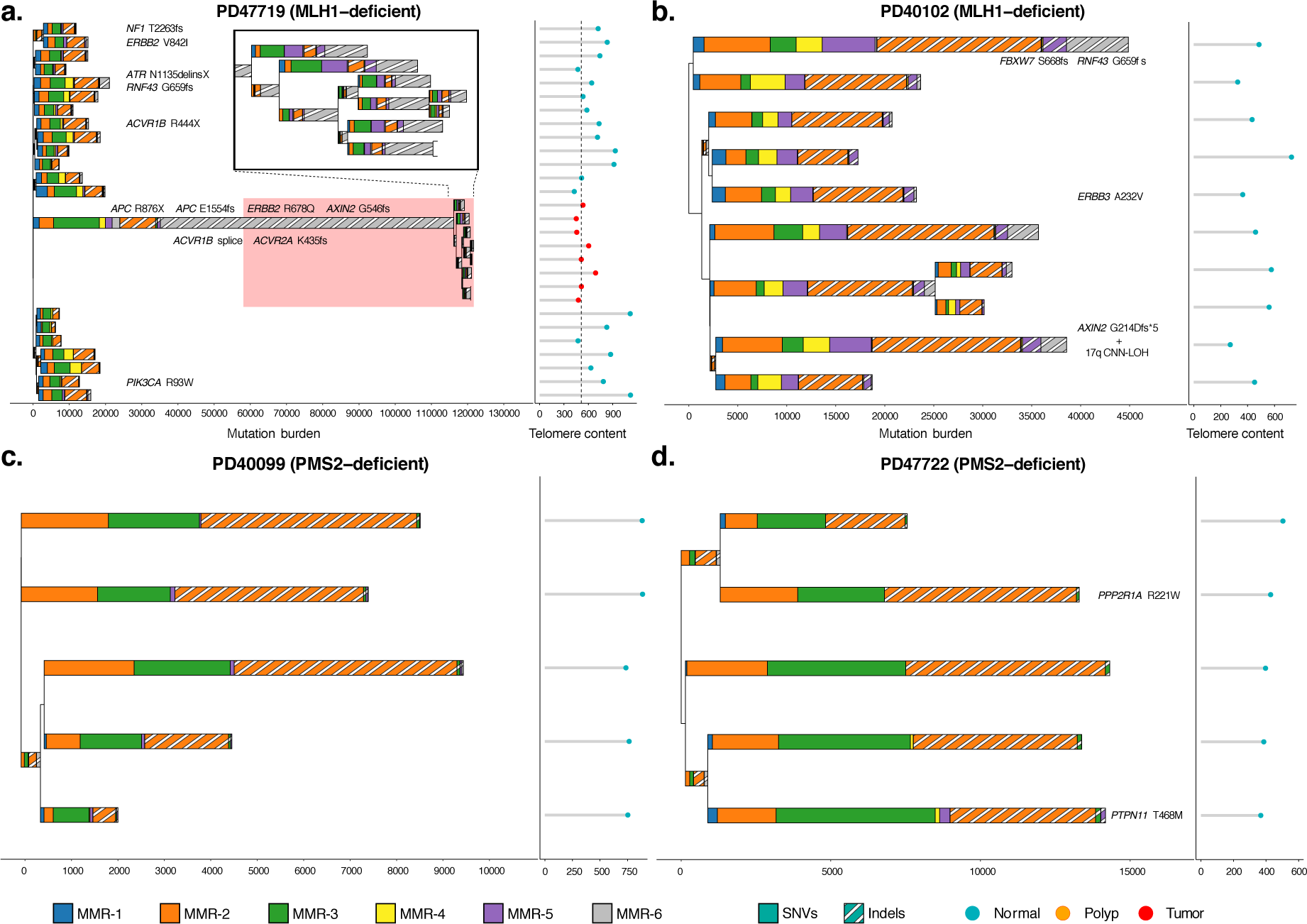
Phylogenetic Mutational Signature Attribution reveals Differential DNA Damage Repair Deficiency in Healthy and Neoplastic MutLα-deficient Intestinal Crypts. (**a-d**) Intestinal crypt phylogenies were inferred from all mutations. Branch length is proportional to the number of mutations detected. Branching indicates a historical crypt fission event. Stacked bar plots represent the mutational signature composition contributing to individual branches and colour-coded according to exposure. Branches defined by <100 mutations were omitted from signature extraction. Stacked bar plot constituents: solid pattern – SNVs, stripped pattern – indels. Right panel: telomere content for individual crypts coloured by disease state. Dashed line indicates the median telomere content for neoplastic intestinal crypts. Coloured rectangular areas highlight branches formed by intestinal crypts microdissected from neoplastic tissues. Detected driver mutations are listed next to or along branches to indicate their presence among related intestinal crypts. **(a)** PD47719 (MLH1-deficient), phylogeny of normal (n = 20) and neoplastic (n = 7) intestinal crypts. Coloured area indicates branches common matched microdissected colon cancer. Inset illustrates branches established in the final stages of colon cancer development. (**b**) PD40102 (MLH1-deficient), phylogeny of only normal (n = 10) intestinal crypts. (**c**) PD40099 (PMS2-deficient), phylogeny of only normal (n = 5) intestinal crypts. (**d**) PD47722 (PMS-deficient), phylogeny of only normal (n = 5) intestinal crypts.

Investigation of potential driver mutations revealed known oncogenic mutations in *FBXW7*, *ERBB2*, *ERBB3*, *PIK3CA*, *TP53*, *PPP2R1A, ACVR1B, ACVR2A, ATR, AXIN2* and *BAP1* in normal intestinal epithelium (**Figure 4**, **Figure S3a**, **Table S5**). Crypts from polyps and tumors showed the expected complement of *APC* mutations (**Figure S3a**). For all MutSα-deficient neoplastic tissues, *APC* mutations were C>T mutations at <u>C</u>G dinucleotides, indicating that MMR-1 can induce driver mutations. Analysis of the driver mutation landscape across all intestinal crypts indicates that MMR-1 is the primary force of driver mutation acquisition, further complemented by microsatellite instability (**Figure S3b**). While MMR-proficient intestinal epithelial cells rarely acquire driver mutations, constitutionally MMR-deficient cells acquire more mutations, at younger ages, especially in cases of MutSα-deficiency (**Figure S3c**). This confirms that MMR-1 represents a major force of early driver acquisition in MutSα-deficient cells.

### Activity of Mutational Processes in Other Normal Tissues

In addition to intestinal epithelium, we microdissected and sequenced 25 samples from 5 CMMRD donors across a wide array of tissues types, including brain cortex, lymphocyte aggregates, intraepithelial nerves and arterial walls. All MMRD genotypes, with the exception of *MLH1*, were represented. Sufficient numbers of SBS and indels were detected for mutational signature extraction (**Figure 5a**, **Table S6**). While mutation burdens varied considerably across the samples based on tissue type and level of clonality, we found that MutSα-deficient donors were characterised by high levels of MMR-1, the 5mC deamination signature, across different tissues; furthermore, this signature was again largely absent from MutLα-deficient samples (**Figure 5b**). This was especially evident in the brain cortex and striatopallidal fibers of the *MSH6*-deficient case PD40100 and to a lesser degree in the brain blood vessel of the *MSH2*-deficient case PD40098 (**Figure 5c**). The only exceptions were large intra-epithelial lymphoid populations in PD40103 (**Figure 5c**), which were enriched for the replication-associated mutational signature MMR-3. The clonality of these samples suggests that single B cells seeded the colonic intra-epithelial and expanded *in situ* to form the observed secondary lymphoid structures, which may explain why replication-mediated signatures predominated. Similar to the colonic epithelium, the tissue samples from MutLα-deficient cases revealed strong contributions of signatures MMR-2 and MMR-3.

**Figure 5.**
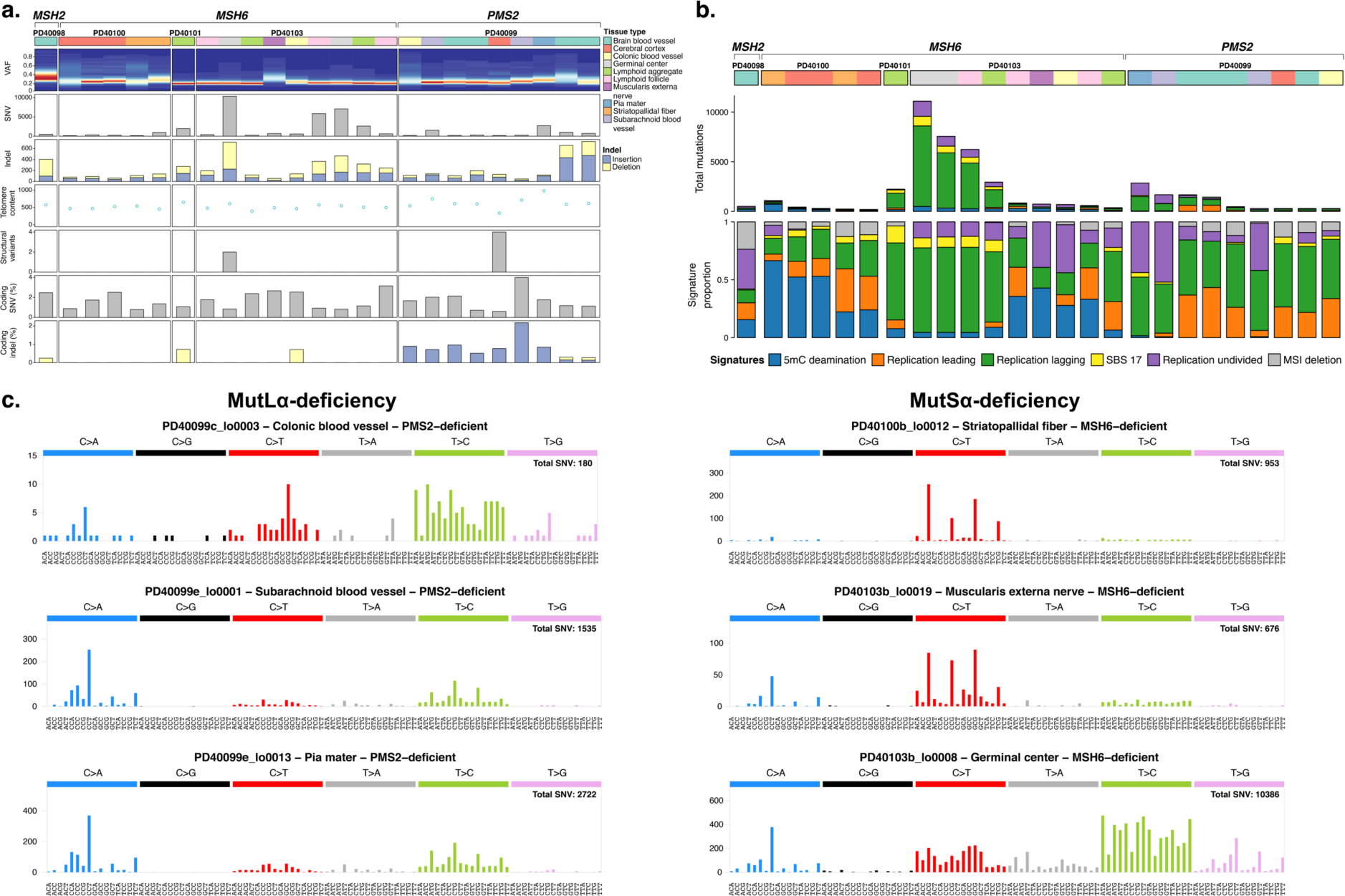
Mutational Signature Analysis of Other CMMRD Tissues Confirms DNA Damage Repair Specificity Among MMR Components. **(a)** Each column indicates an individual sample. From top to bottom: tissue type indicator, VAF distribution heatmap, SNV burden, indel burden distinguished by insertions and deletions, telomere content, SV burden, CNV burden, percentage of coding SNVs, percentage of coding indels distinguished by insertions and deletions. **(b)** Total and relative mutational signature contribution for microdissected non-intestinal samples as identified by the indicated colour-coding. Legend is shared with panel (**a**). **(c)** Left: typical mutational spectra for different MutLα-deficient tissues. Right: typical mutational spectra for different MutSα-deficient tissues.

Taken together, these data indicate that while MMRD genotype imposes a strong influence on the composition of mutational signatures across tissues, there are also detectable differences based on tissue type that could be the consequence of stem cell dynamics or environmental factors. This suggests that tissue-specific DNA damage influences the effects of MMRD on the observed mutational landscape.

### Distribution of Mutations Across the Genome

The MMR pathway is known to preferentially correct mutations in early replicating regions and even more efficiently in exons, meaning that its deficiency equalizes mutation rates across the genome (Frigola et al., 2017; Supek and Lehner, 2015). In our data, we found that this equalization of mutation rates by replication timing was more pronounced in cases of MutSα-deficiency (loss of either *MSH2* or *MSH6*) than MutLα-deficiency (loss of either *MLH1* or *PMS2*; **Figure S4a-b**). This difference was especially apparent in exonic regions, where MutSα-deficient cases acquired a median 50 exonic substitutions per year compared to a median 19 exonic substitutions per year for cases of MutLα deficiency. Thus, our data demonstrate that the targeting of mismatch repair to early replicating and exonic regions is largely dependent on MutSα rather than MutLα.

Across the genome, we observed that broad domains of late-replicating DNA had high relative mutation rates in cases PD40102, PD47719 and PD47722, all MutLα-deficient patients; a more pronounced variation than seen for MutSα-deficient patients (**Figure 6a-b**). In contrast, cases of MutSα-deficiency exhibit increased relative mutation rates at regions of high CpG density, presumably reflecting the greater opportunity for 5mC deamination in such regions (**Figure 6b**, **Figure S4c**). Indeed, using colonic whole genome methylation data, we found a strong linear relationship between the density of methylated CpG dinucleotides and the numbers of C>T mutations in a <u>C</u>G context (**Figure 6c**). Correcting the relative mutation rates of <u>C</u>G><u>T</u>G for overall CpG density only partially removed differences across genomic elements; however, correcting for methylated CpG density completely eliminated these differences (**Figure 6d**).

**Figure 6.**
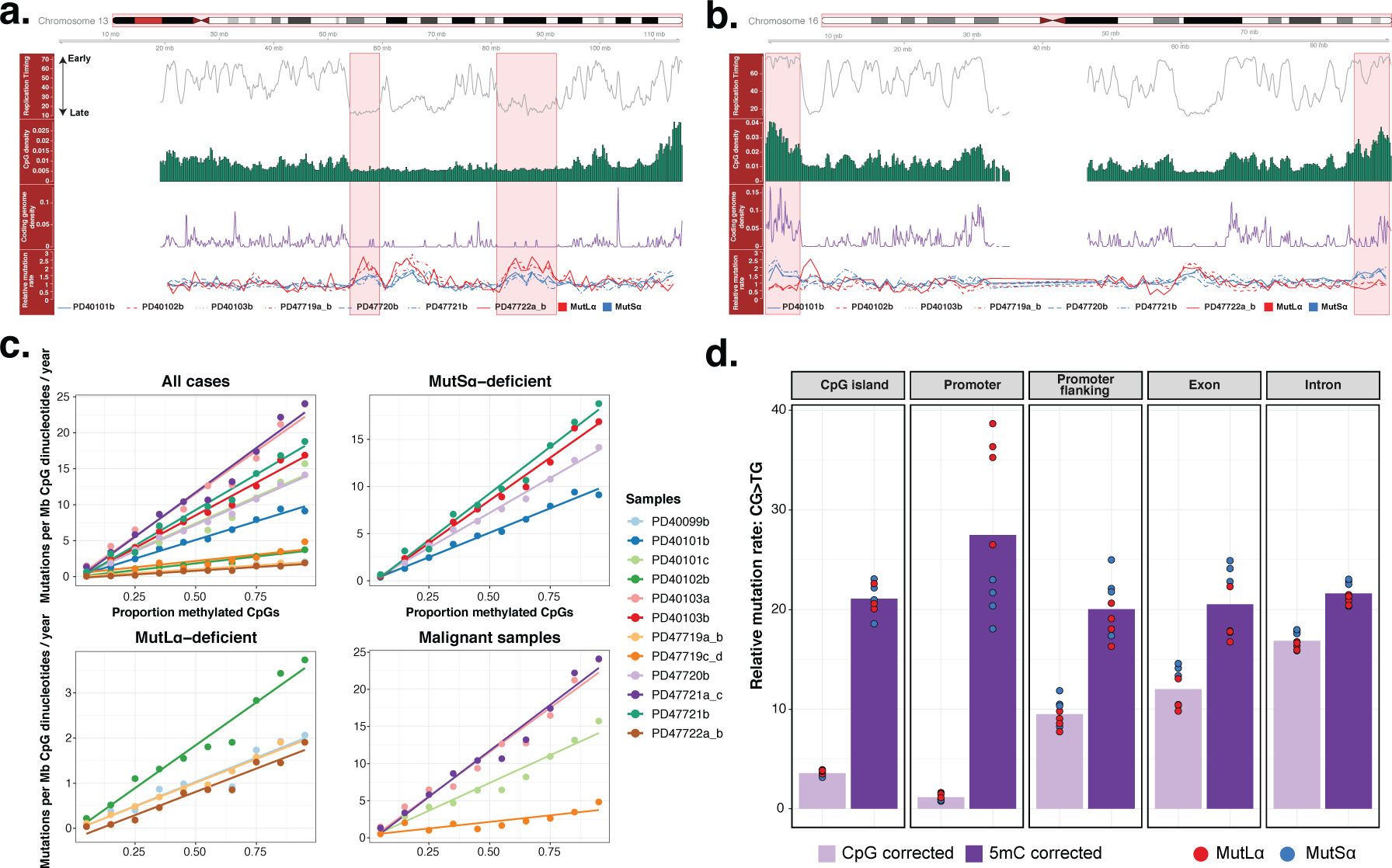
Determinants and Genomic Distribution of DNA Damage Typically Repaired by MMR. **(a)** Distribution of determinants and mutations on chromosome 13 across non-overlapping 1 Mb bins. From top to bottom. (I) Average replication timing profile. High values indicates early replication and vice versa. (II) CpG-dinucleotide frequency. (III) Coding genome frequency. (IV) Average relative SNV rate of healthy intestinal crypts across the non-overlapping bins. Line type indicates individual donors whereas line colour indicates which MMR complex is deficient. Highlighted areas indicate broad domains of late genome replication. Case PD40099 was excluded on the basis of low genome-wide mutation counts. **(b)** Distribution of determinants and mutations on chromosome 16 across non-overlapping 1 Mb bins. Information included and colour coding is the same as panel (**a**). Highlighted areas indicate areas of high CpG and coding genome density. **(c)** Linear relationship between methylation levels and methylation damage. CpG-dinucleotides are grouped on observed methylation levels (methylation level bins) detected in a normal sigmoid colon. Mutation burden was determined across all bins and corrected for total CpG counts in millions and age. Solid line is the results of ordinary linear regression. Data for all microdissected healthy intestinal crypts and deficiency of individual MMR complexes is displayed in the top row and lower-left panel. Data for neoplastic samples are displayed in the lower-right panel. **(d)** Relative mutation rates (RMR) of methylation damage (<u>C</u>G><u>T</u>G) across different genomic elements further distinguished by MMR complex deficiency. RMR values are obtained per genomic element by correcting the total <u>C</u>G><u>T</u>G burden per sample by either CpG density or methylation levels.

Regions with high CpG density tend to be early replicating and more gene-dense, suggesting that some of the apparent equalization of mutation density in MutSα-deficient cases could be caused by balanced effects of replication-dependent signatures MMR-2 and MMR-3 in late-replicating regions versus MMR-1 in early-replicating regions. We therefore studied the distribution of individual MMR signatures across the genome. In MutSα-deficient intestinal epithelium, MMR-1 contributes to >80% of somatic mutations in early-replicating regions (**Figure S5**, **Table S7**) and >90% in coding sequence (CDS) (**Figure S6**, **Table S8**), whereas replication-dependent mutational processes are more evident in later-replicating regions or outside the coding genome. Mutations arising from MMR-1 in the most functional genomic regions is much more a feature of MutSα-deficiency than MutLα-deficiency (for earliest-replicating genome: effect size 4.25; CI_95_%=1.94-6.51; *P*=0.01; for CDS: effect size 2.98; CI_95_%=1.52-4.41; *P*=0.01; generalized linear mixed models comparing MutSα-deficient and MutLα-deficient samples).

In summary, then, mutations arising from deamination of methylated cytosine (MMR-1) concentrate in the most functional coding and early replicating regions of the genome, whereas mutations arising from replication errors (MMR-2, MMR-3, MMR-5) predominantly distribute to late-replicating regions of the genome. In the absence of MutSα, these distributions balance almost exactly, leading to broadly equal mutation rates across the genome, whereas in MutLα-deficiency, the replication-associated signatures are predominant such that mutation rates are higher in late-replicating regions.

### Verification of Genotype-specific Mutational Signatures in Sporadic Cancers

Our investigation into subjects born without mismatch repair reveals that the mutational consequences are shaped by MMRD genotype, tissue type and neoplastic transformation. To assess whether these observations generalize, we analysed sequencing data from sporadic MMRD tumors, collated from TCGA, CMMRD cancers and cancers sequenced in-house. The germline and somatic landscape were screened for substitutions, indels, SVs and CNVs that perturb any of the 4 genes encoding the core MMR pathway. Cases with disruptive MMR mutations (nonsense, frameshift, splice site, copy number loss or structural variation) present as homozygous, compound heterozygous variants or affecting two MMR genes were identified. For missense substitutions, only clinically relevant variants recorded in ClinVar, InSiGHT or the literature were classified as pathogenic. Cases with documented MMRD or those with high mutation burdens and at least one MMRD-associated mutational signature were selected for further characterization.

With this approach, we identified 117 cases with an established MMRD genotype, spanning endometrial cancer, colorectal adenocarcinoma and gastric cancer most frequently, but also occasional cases of prostrate adenocarcinoma, breast cancer, brain cancers, sarcomas and other cancers (**Figure 7a**, **Table S9**). Sporadic MMRD cancers were most often defined by the loss of a single MMR component, although disruptive mutations affecting two MMR genes were also seen, with *MSH2* and *MSH6* being the most common co-mutated pair (**Figure 7b**). We used Dirichlet regression on the mutational signature composition to infer associations between mutational signature exposure and MMRD genotype. *PMS2*-deficient cases were used as reference. Replicating the observation made in congenital cases, activity of signature MMR-1, reflecting methylated cytosine deamination, was significantly greater in tumors with loss of *MSH2* (*P*=1.66x10^-5^) or *MSH6* (*P*=1.64x10^-5^; **Figure 7c**) (**Table S10**). Exposure to MMR-5, found in MutSα and *MLH1*-deficient CMMRD tissues especially upon neoplastic change, was similarly detected in sporadic MMRD cancers with these genotypes (*MLH1*: *P*=1.3x10^-3^; *MSH2*: *P*=1.34x10^-4^; *MSH6*: *P*=0.02; **Figure 7c**, **Table S10**). Comparing MutSα-deficient to MutLα-deficient cancers revealed that the association between MutSα loss and increased MMR-1 activity could be replicated across different cancer subtypes (all, endometrial, colorectal and rare MMRD cancers; **Figure S7a**, **Table S10**). The association between MutSα loss and MMR-5 activity was confirmed in endometrial and colorectal MMRD cancers (**Figure S7b**, **Table S10**).

**Figure 7.**
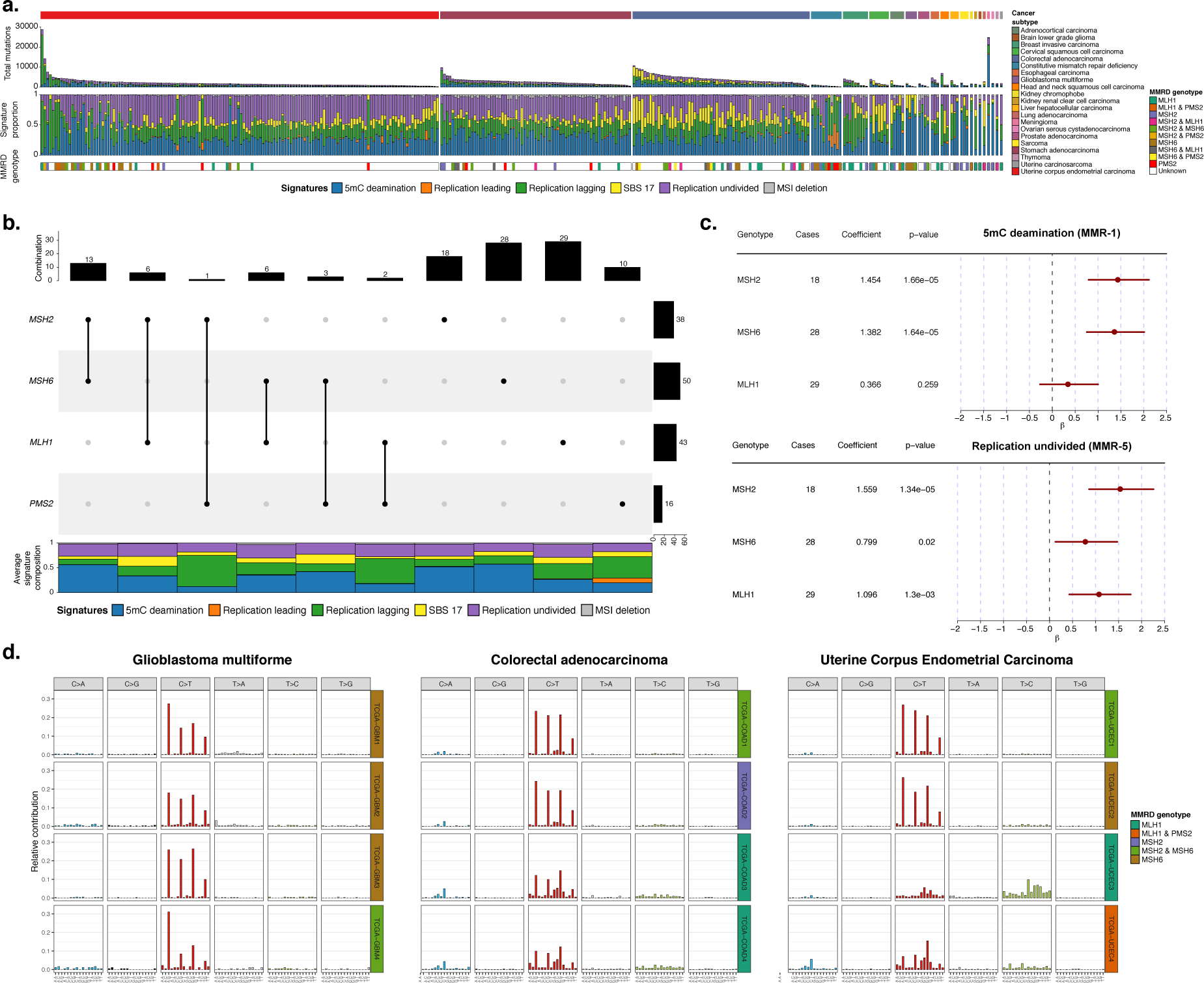
MutSα is a Primary Repair Force Engaging Methylation Damage as Confirmed in MutSα-deficient Cancers. **(a-e)** MMRD status was determined across all TCGA cancer subtypes and supplemented by a CMMRD WES cancer cohort. MMRD genotype was determined by screening for loss-of-function or known damaging variants, CNVs and SVs involving MMR components. Cases with MMRD as a consequence of MMR component promoter silencing were listed as unknown. **(a)** Mutation burden, mutational signature contribution and MMRD genotype for MMRD cancers present in the combined cohort. Each column represents an individual MMRD cancer. From top to bottom: cancer subtype, mutational signature attribution based on all mutations (SNVs and indels), relative mutational signature composition and MMRD genotype indicator. A white MMRD genotype indicator indicates either an unknown genotype or that MMRD is established through MMR gene promoter silencing. **(b)** UpSet plot summarizing MMRD genotype counts across all genotyped MMRD cancers. Top: number of cases with the indicated MMRD genotype combination. Right: number of cases per MMR component lost. Bottom: average mutational signature contribution per indicated MMRD genotype. **(c)** Forest plots summarizing the Dirichlet regression results. PMS2-deficient cancers were used as reference. Cancers with components lost from both MMR complexes (MutSα and MutLα) were excluded. Significant associations between MMR component loss and mutational signature activity was seen for the MMR-1 and MMR-5 signatures. **(d)** SBS mutational profiles per indicated cancer subtypes across different MMRD genotypes. Coloured label on the right of each mutational profile indicates the established MMRD genotype. Left: mutational profiles of MMRD glioblastoma multiforme enriched for loss of MutSα. Middle: colorectal adenocarcinoma (COAD), top 2 rows are typical mutational profiles for MutSα-deficient cases while the bottom 2 rows are indicative for MutLα-deficient cases. Right: endometrial cancer, top 2 rows are typical mutational profiles for MutSα-deficient cases while the bottom 2 rows are indicative for MutLα-deficient cases.

As previously reported (Touat et al., 2020), defects in *MSH2* and *MSH6* were strongly associated with glioblastoma multiforme in our dataset. As a result, the mutational spectrum of these tumors was dominated by C>T mutations at <u>C</u>G dinucleotides (**Figure 7d**). For colorectal and endometrial cancers, cases included both MutSα-deficient and MutLα-deficient genotypes, and the mutation spectrum differed accordingly. Overall, then, the differences across genotype and tissue type in CMMRD individuals generalize to sporadic MMR deficiency.

## Discussion

The mismatch repair pathway has long been recognised as a major bulwark in the defence of our genome – our data now quantify the magnitude of its activity in the day-to-day life of humans. We see rates of 500-3000 base substitutions and indels per year per cell in patients with deficiency of a single component of the pathway, compared to ∼50/year in normal individuals (Lee-Six et al., 2019). Furthermore, we observe considerable differences in mutational signatures between individuals inheriting deficiency of MutSα and those without MutLα. Together, these observations suggest that each component of the MMR pathway normally repairs 1-10 potentially mutagenic events every day in every intestinal cell, on average, and the two major components target different forms of DNA damage or replication error. This is in normal cells – once a clone begins to exhibit neoplastic growth, MMR becomes an even more critical defence, likely repairing tens of thousands of additional events over short periods of time.

Mismatch repair is classically described as a replication-linked process – errors introduced by replication polymerases, both substitutions and indels, are recognised by MutSα, triggering recruitment of MutLα, excision of the mismatch from the nascent strand and repair by replication from the template strand (Kunkel and Erie, 2015). In our data, the MMR-2 and MMR-3 signatures likely capture this canonical replication-linked repair – the two signatures are found in both MutSα and MutLα deficiency, show clear biases by leading or lagging strand of replication respectively, and vary in intensity across the genome by replication timing. The two signatures reflect the different patterns of replication errors made by leading- and lagging-strand polymerases, with small insertions dominating the leading-strand signature (MMR-2) and T>C transitions dominating the lagging-strand signature (MMR-3). Beyond MutSα, it is known that MutSβ, comprising MSH2 and MSH3, also signals to MutLα to repair errors introduced during replication, especially deletions (Romanova and Crouse, 2013). We find that the MutSβ pathway is numerically more important as the MutSα pathway for preventing indels in normal human cells, evidenced by the observation that the rate of indels in deficiency of *MSH6*, not a component of MutSβ, is a quarter of the rate seen with deficiency of *MSH2*, present in both MutSα and MutSβ, or with deficiencies of *PMS2* and *MLH1*, components of MutLα (**Figure 1c**). The MMR-6 signature in part captures this MutSβ repair process, as it has considerably lower contribution to *MSH6*-deficient cases than *MSH2* deficiency, and is dominated by deletions.

An unexpected finding is that MutSα is the major repair pathway in human cells against spontaneous deamination of methylated cytosine to thymine, a repair activity that cannot be linked to replication. Our evidence for this is that signature MMR-1 is numerically the dominant process in MutSα deficiency, a signature comprising almost purely C>T substitutions at CpG dinucleotides, with no replication strand or transcription bias, correlating linearly with the fraction of methylated cytosines across the genome. Deamination of methylated cytosine is a spontaneous hydrolytic reaction in double-stranded DNA (Shen et al., 1994). As a spontaneous reaction, it will occur in all phases of the cell cycle, including interphase. Because the deamination generates a thymine, if the G:dT mismatch is not repaired before replication, the event will become fixed as a somatic mutation – hence, this repair activity must occur before replication. Interestingly, we find no increase in the rates of this mutational signature in MutSα-deficient cells after neoplastic transformation, consistent with it being independent of replication. Furthermore, the tumor types with sporadic MMR deficiency showing the highest rates of MMR-1 are those with low intrinsic cell division rates, tissues such as meninges, pleura and glial cells (**Figure 7**).

In MutSα-deficient intestinal crypts, 350-1150 C>T substitutions at CpG dinucleotides accrued every year. Spontaneous deamination of methylated cytosine is estimated to occur at rates of 5.8x10^-13^/5mC/second at 37°C (Shen et al., 1994) – with methylated cytosines comprising ∼1% of the human genome (Ehrlich et al., 1982), this rate equates to ∼1000 deamination events per diploid genome per year. The striking correspondence of the expected number of spontaneous deamination events with our observed mutation rates argues that MutSα is the critical component of our defence against this mutational process. The paucity of MMR-1 in patients with *MLH1* or *PMS2* deficiency implies that MutLα is not the partner for repair of these deamination events. Instead, a strong candidate is *MBD4*, a thymine DNA glycosylase that binds G:T mismatches in a methylated cytosine context (Hendrich et al., 1999). Germline deficiency of *MBD4* causes tumors with remarkably elevated C>T mutation rates at CpG dinucleotides, with mutation rates and mutation spectrum very similar to that of MMR-1 here (Derrien et al., 2020; Pan-cancer Analysis of Whole Genomes Consortium, 2020; Sanders et al., 2018).

These data demonstrate that studying somatic mutations in normal tissues from natural human knock-outs can enrich observations from experimental model systems and cancers. By comparing mutation rates to normal tissues in healthy controls, we have quantified the day-to-day activity of MMR in normal human cells across tissue types; by comparing mutational signatures and rates across genotypes, we have separated the relative contributions of individual components of the MMR pathway to different types of DNA lesions; by comparing normal and transformed cells from the same patients, we have shown the increased dependence on MMR’s protective activity caused by rapid cellular proliferation.

## ACKNOWLEDGMENTS

We thank the staff of the Wellcome Sanger Institute Sample Logistics, Sequencing and Informatics facilities for their contribution, including Laura O’Neill, Calli Latimer and Kirsty Roberts for their support with sample management; Remco Hoogenboezem and Peter Valk for help with analyses, advice and discussions; Stefan Dentro for help with analyses; and Iñigo Martincorena for advice and discussions. Results presented here are in part based upon data generated by the TCGA Research Network: https://www.cancer.gov/tcga.

## AUTHOR CONTRIBUTIONS

M.A.S., U.T. and P.J.C. conceived the study design. V.F., B.B.C., M.E., V.B., A.S. and U.T. recruited individuals, collected samples, and curated clinical data. L.M. and M.A.S. developed bespoke DNA library preparation methods. L.M., Y.H. and T.B. helped with sample preparation, tissue fixation, tissue sectioning and sample submission. H.L. and M.R.S. contributed and analysed control data. C.F., R.B. and I.J.M. provided help and advice on analysing TCGA data. U.M. provided sequencing data. P.S.R. and T.H.H.C. provided a bespoke procedure for unmatched variants filtering. H.S.V. and M.G. performed signature extraction and fitting. M.A.S., H.S.V., M.G. and P.J.C. performed data analysis. P.J.C. and M.G. oversaw statistical analyses. P.J.C. and M.A.S. drafted the manuscript, with input from V.F. and U.T.

## CODE AVAILABILITY

Code required to reproduce the analyses in this paper are available online (https://github.com/MathijsSanders/MMR). Mutation calling algorithms are available through GitHub (https://github.com/cancerit). All other code is available from the authors upon request.

## DATA AVAILABILITY

Whole genome and exome sequencing BAM files for cases reported in this study have been deposited in the European Genome-Phenome Archive (EGA) under accession code EGAS00001002881 (https://www.ebi.ac.uk/ega/home).

## METHODS

### Human Tissue Samples

All human biological samples were collected with informed consent by the Biallelic Mismatch Repair Deficiency Syndrome (bMMRD) biobank located at The Hospital of Sick Kids, Toronto, Canada, in accordance with procedures approved by the Local Research Ethics Committee (REC reference 17/NW/0713). Samples were snap-frozen in liquid nitrogen or stored at -80 °C upon collection and stored at -80 °C after transfer to the bMMRD biobank at The Hospital of Sick Kids.

### Tissue Preparation

Frozen tissues were first thawed at 4 °C for 10-15 min. Larger biopsies were dissected into smaller sections which were stored at -80 °C to safeguard materials for future experiments. Thawed tissues were fixed in 70% ethanol and embedded in paraffin by the Tissue-Tek VIP 6 AI tissue processing instrument (Sakure) based on an ethanol-only embedding protocol. Sections of 10μm thickness were cut from the paraffine-embedded tissue blocks and mounted onto poly-ethylene naphtholate (PEN)-membrane slides (Leica). Tissue slides were stained with the following haematoxylin and eosin (H&E) protocol in sequential steps: xylene (2 min, twice), ethanol (100%, 1 min, twice), deionized water (1 min, once), Gill’s haematoxylin (10-20 s, once), tap water (20 s, twice), eosin (10 s, once), tap water (10-20 s, once), ethanol (70%, 20 s, twice), and xylene (10-20 s, twice). Tissue slides were cover-slipped after H&E staining for scanning with the NanoZoomer S60 Slide Scanner (Hamamatsu), generated images were viewed with NDP.View2 (Hamamatsu).

### Laser Capture Microdissection

Areas of interest were identified from the scanned slides. Laser capture microdissection of these areas was performed with the LMD7000 microscope (Leica) and collected into a skirted 96-well PCR plate. Overview pre- and post-dissection images were stored for quality control. For some donors bulk samples from connective or muscle tissues were taken to be used as matched control. Cell lysis was achieved using 20μl of proteinase-K Picopure DNA Extraction kit (Arcturus). Thermocycler protocol: sample incubation at 65 °C for 3 hours followed by proteinase denaturation at 75 °C for 30 minutes. Processed samples were stored at -20 °C prior to submission for DNA library preparation.

### Low-input DNA Library Preparation and Sequencing

DNA library preparation was undertaken as previously described using a low-input DNA library preparation method (Ellis et al., 2021). This protocol allows for high quality DNA library preparation starting from low material quantities without whole genome amplification. DNA library concentration was assessed after library preparation, a minimum library concentration of 5 ng/μl was used to determine whether to take DNA sequencing forward. All LCM samples were subjected to 2x150bp paired-end whole genome sequencing. Samples from donors PD40098, PD40099, PD40100, PD40101, PD40102 and PD40103 were sequenced on the HiSeq X platform (Illumina), while samples from donors PD47719, PD47720, PD47721 and PD47722 were sequenced on the NovaSeq 6000 platform (Illumina) with the XP kit and using unique dual indexes. Pools of samples were sequenced to achieve a coverage of *≥* 30.

### SBS Calling

Sequencing data were aligned to the NCBI human reference genome (GRCh37) using the Burrow-Wheeler Aligner (BWA-MEM) (Li and Durbin, 2009). Duplicated mate-pairs were marked for removal during downstream analysis. Single Base Substitutions (SBS) were called using ‘Cancer Variants through Expectation Maximization’ (CaVEMan) (Jones et al., 2016). Mutations were called against a synthetic unmatched normal control. Post-calling filters were utilized to remove low-quality variants, alignment errors, low-input DNA library preparation specific artefacts and germline variants as previously described (Lee-Six et al., 2019; Moore et al., 2020). The following filters were applied: (1) common single nucleotide polymorphisms and recurrent artefacts were removed by filtering against a panel of 75 unmatched normal samples (Nik-Zainal et al., 2012), (2) common artefacts and erroneous variants as a consequence of poor alignment were removed by applying a threshold for the median alignment score (ASMD *≥* 140) of reads supporting variants and less than half of the variant-supporting reads could be clipped (CLPM = 0), (3) by counting variant supporting fragments instead of reads to prevent counting variant-supporting mate-pairs double and (4) a filter that detects variants introduced due to incorrect processing of cruciform DNA during DNA library preparation. Software implementing these filtering steps can be found at https://github.com/MathijsSanders/SangerLCMFiltering.

A subsequent set of filters is applied to remove retained germline variants and potential artefacts. Code for performing these filtering steps can be found at https://github.com/TimCoorens/Unmatched_NormSeq. Mutations across all intestinal samples were aggregated and read pile-up was performed with AnnotateBAMStatistics (https://github.com/MathijsSanders/AnnotateBAMStatistics) to determine the fragment counts of the mutant and wildtype alleles per mutation. Germline variants were filtered using an exact binomial test. Per donor mutant counts were aggregated across all intestinal samples in conjunction with the total fragment coverage. Taking the fragment variant and coverage counts a variant is retained when it deviates significantly from the expected values from a binomial distribution with p = 0.5. A second filter is used to filter potential remnant artefacts present at lower variant allele frequencies (VAF). For somatic variants a significant overdispersion is present when fitting a beta-binomial distribution. Variants consistently present with similar VAFs across all included samples are likely erroneous and yield little overdispersion. Therefore, a beta-binomial test was applied. Genuine variants were determined by using a overdispersion threshold (rho) of *≥* 0.1. This process yielded the final list of somatic variants used for downstream analysis.

### Indel Calling

Indels were called by cgpPindel (Raine et al., 2015) using a synthetic normal, described under ‘SBS calling’, as unmatched control. Indels passing the standard filtering were subjected to a quality score *≥* 300 and should be at least covered by 15 reads. Detected indels positioned in large repeats (> 9bp) are flagged by cgpPindel and removed from the final list of somatic variants. The number of mutant reads, wildtype reads and VAF was determined by cgpVAF. After collating these statistics, a similar protocol for determining germline variants and possible artefacts was used as described for SBS.

### Deletion-to-indel Ratio

The deletion:indel ratio was determined for each normal intestinal crypt sequenced to assess the linear relationship between these lesions dependent on the mismatch repair deficiency (MMRD) genotype. A linear mixed model was constructed with lme4 (Bates et al., 2014) to model the number of deletions as a dependent variable on the interaction effect insertions:genotype (the number of insertions and MMRD genotype, fixed effect), donor label (random intercept to account for multiple samples per donor) and genotype-dependent insertions (random slope for insertions based on MMRD genotype). Crypts from neoplastic intestinal tissues have a strong phylogenetic relationship resulting in similar numbers of indels. The average deletion:indel ratio was calculated for these samples as an indication of which indel lesion type are often acquired.

### Structural Variant Calling

Structural variants (SV) were called using a matched, but phylogenetically unrelated, sample from the same donor with the ‘Breakpoints Via Assembly’ (BRASS) algorithm (Campbell et al., 2008), with further annotation from GRASS (https://github.com/cancerit/BRASS). SVs were annotated further by AnnotateBRASS (https://github.com/MathijsSanders/AnnotateBRASS) as previously described in detail (Moore et al., 2020). In brief, low-input DNA preparation results in the erroneous introduction of SV at low read counts. AnnotateBRASS determines the following per SV: the number of supporting read pairs, the variance in the alignment position of read pairs, whether read-pairs are clipped or carry an excess of variants not reported in single nucleotide polymorphism databases, are in the correct orientation or whether SV-supporting read-pairs are in regions marked by high proportions of other read-pairs aligning to different parts of the genome (high homology). After annotation SVs are filtered as previously described in detail https://github.com/MathijsSanders/AnnotateBRASS (Moore et al., 2020).

### Copy Number Variation Calling

Coverage and B-allele frequency (BAF) information was extracted for samples included in the study by ConstructASCATFiles (https://github.com/MathijsSanders/ConstructASCATFiles). Single nucleotide polypmorphisms (SNP) with a population frequency greater than 0.01 were used for extracting coverage and BAF statistics across all samples. Coverage and BAF statistics were grouped by donor and assessed for quality via ‘QualityControl_and_PCA.R’ (https://github.com/MathijsSanders/PREASCAT). For each donor one or more control samples are defined which are assumed to comprise fully diploid cells. This could either be bulk polyclonal samples (connective tissue) or other tissues for which it is certain that the genome is diploid. SNPs with limited coverage across all control samples are excluded from the analysis. Samples are corrected for library size by assuming that heterozygous SNPs with a balanced VAF (∼ 0.5) represent diploid genomic areas. The LogR ratio is determined by dividing the library-size corrected coverage for each samples with the median library-size corrected coverage of the controls. The X chromosome is multiplied by 2 for male individuals to correct for sex differences. The low-input DNA library preparation protocol on occasion results in substantial variance in the BAF of heterozygous SNPs in the control samples. The median BAF across the control samples is used to determine the set of heterozygous SNPs for each donor. Principal component analysis (PCA) was applied to normalized LogR values to detect systemic biases present in all samples or the samples of the donor of interest. Among such areas are the HLA locus and peri-centromeric regions and regions of high-vs-low CpG density. Final LogR and BAF statistics are calculated with ‘construct_ASCAT_file.R’ (https://github.com/MathijsSanders/ConstructASCATFiles). This algorithm follows the same approach but includes prinicipal component (PC) regression to assuage any systemic noise present. LogR profiles are centered around 0 by subtracting the mean value followed by PC regression to determine the absolute presence of noise. This fit is extracted from the 0-centered LogR profile followed by restoration to the previous average value by adding the mean value earlier used for subtraction. Finally, the resulting files are used in ASCAT for allele-specific CNV detection (Van Loo et al., 2010).

### Telomere Content Estimation

Attrition of the telomeres is a hallmark of cellular aging. Assessment of telomere lengths for all microdissected samples, including neoplastic intestinal epithelium, was achieved by two distinct approaches.

Telomerecat v3.4.0 (Farmery et al., 2018) was used for estimating telomere lengths based on WGS data with length correction enabled and the number of simulations set to 100. In rare instances telomere lengths could not be estimated. One possible explanation is the use of the HiSeq X and NovaSeq 6000 instruments for distinct sets of microdissected samples. A second telomere length algorithm, TelomereHunter (Feuerbach et al., 2019), was used as an orthogonal approach which yielded results for all samples. Strong correlation was observed upon comparing the different telomere length estimation algorithms. Telomeres from neoplastic intestinal epithelium were significant shorter compared to their synchronously collected matched normal tissues indicative that neoplastic change results in increased proliferation rates and shorter telomeres. Telomere content estimates from TelomereHunter were reported because information was available for all samples.

### Inference of Phylogenetic Trees

Phylogenies from all mutations, SBS and indels, of normal and neoplastic colonic crypts (n=110) was reconstructed for 8 CMMRD donors across all MMRD genotypes. SBS were called by CaVEMan as described under ‘SBS calling’, while indels were called by cgpPindel as described under ‘Indel calling’. Finally mutations were recalled across all intestinal crypts samples for each donor using AnnotateBAMStatistics (SBS) or cgpVAF (indel). Variants with a VAF *≥* 0.3 were labelled as present (1), VAF *≤* 0.1 as absent (0) and a VAF between 0.1 and 0.3 (?) as ambiguous. This string of labels was used as input for phylogenetic tree reconstruction using a parsimonious approach in MPBoot (Hoang et al., 2018), bootstrap parameter set 1.000, and mutations were mapped to branches of the phylogenic tree using maximum likelihood estimation.

### Mutational Signature Extraction

All classes of SBS, multi-nucleotide variants (MNV) and indels were used for mutational signature extraction. For normal and neoplastic intestinal crypts branches from the reconstructed phylogenetic trees were used as input rather than the complete per-sample mutational catalogue. Some intestinal crypts have strong phylogenetic relationship due to crypt fission at a later point in life. Using branches from the phylogenetic trees prevents mutations being duplicated in the same mutational signature extraction procedure. Additionally, using the branches as input gives insight into the changes of the mutational signature composition upon neoplastic change. The full mutational catalogue for non-intestinal epithelium microdissected samples is used due to little genetic kinship to other non-intestinal samples from the same tissue. Mutational signature analysis was performed using TensorSignatures (Vöhringer et al., 2020), which defines mutational signatures in terms of SBS, MNVs as well as indels. SBS are counted strand symmetrically (C>A, C>G, C>T, T>A, T>C and T>G) and were classified based on the 5’ and 3’ flanking nucleotide context. MVS and indels were classified by variant type and increasing length. TensorSignatures deconvolves mutational signatures further on the basis of genomic properties by grouping SBS by genomic contexts including transcription and replication orientation, epigenetic and nucleosomal context, and clustering status. Transcription states include the coding, template and unknown strand and were derived from GENCODE v19 annotations. Assignment to replication orientation (leading, lagging or unknown strand) were based on consensus annotations from Repli-seq profiles of GM12818, K564, HeLa, HUVEC and HepG2 cell lines (Hansen et al., 2010). Epigenetic states (TssA, TssAFlnk, TxFlnk, Tx, TxWk, EnhG, Enh, Znf/Rpts, Het, TSSBiv, BivFlnk, EnhBiv, ReprPC, ReprPCWk and Quies) stem from consensus ChromHMM annotations derived from the 127 ENCODE cell lines (Roadmap Epigenetic Consortium). SBS assignment to nucleosomal status (minor grooves facing away from and towards histones, and linker DNA between nucleosomes) were based on MNase cut efficiency data, locating conserved nucleosome dyad (midpoint) positions in human lymphoblastoid cell lines (Pich et al., 2018). Clustered SBS were identified using a 2-state Hidden-Markov model (Vöhringer et al., 2020). The analysis was performed for decomposition ranks ranging between 2-20 and the overdispersion parameters: 10, 20, 30, 50, 100, 1,000 and 10,000. The model with 6 (overdispersion = 10.000) mutational signatures was selected as suggested by the Bayesian information criterion estimator. Optimization of each model was performed for 50.000 training epochs using an ADAM optimizer and an exponential decaying learning rate (starting learning rate 0.1).

### Phylogenetic Tree Visualisation

The mutational composition per phylogenetic tree branches was estimated by TensorSignatures (Vöhringer et al., 2020). The structure of the reconstructed tree was determined as described under ‘Inference of phylogenetic trees’. The fortified tree was plotted by ggtree (Yu, 2020), while the stacked bar plots were drawn upon the tree with ggplot2 and ggpattern (https://github.com/coolbutuseless/ggpattern). Length of branch indicates the number of SBS and indels detected, while the coloured segments of the stacked barplot indicates the contribution of each mutational signature. The SBS and indels were separated for each branch. Coloured solid segments indicate the number of SBS contributed per mutational signature while the coloured striped segments indicate the number of indels contributed per mutational signature. Telomere lengths were added with aplot and ggpubr.

### Driver Mutation Calling

Driver mutation analysis was performed by taking two separate approaches. First, dNdScv (Martincorena et al., 2017) was used to identify mutations under positive selection by comparing the observed nonsynonymous:synonymous mutation ratio to the expected value after correcting for mutational context biases and local genomic information, such as replication timing or local epigenetic state. Mutations were extracted from the phylogenetic trees to prevent variant duplication. Settings were kept at default because none of the samples violated the predefined condition of having a moderate amount of coding mutations or an excess of mutations for a single gene. When including all dissected intestinal crypts, irrespective whether normal or neoplastic, yielded only the gene *APC* as significantly enriched (Benjamini-Hochberg corrected q-value *≤* 0.05) for disruptive mutations which is not unexpected given the inclusion of neoplastic intestinal tissues. For normal tissues none of the recurrently mutated genes were considered significantly enriched for non-silent mutations, even when performing restricted hypothesis testing.

Yet, there were multiple mutations detected in the normal intestinal crypts with amino acid changes often seen in intestinal cancers as reported by COSMIC (Forbes et al., 2008). In the second phase SBS and indels were collated across the normal and neoplastic intestinal crypts separately. Analysis was restricted to protein-coding changes and essential splice acceptor or donor sites. Mutations were annotated with ANNOVAR (Wang et al., 2010), including COSMIC db v91, and compared to a list of 369 driver genes previously published (Martincorena et al., 2017). All types of premature truncating variants (PTV; frameshifts, stopgains or essential splice-site mutations) were considered sufficient for genes known to act as tumour suppressors or have been associated with frequent PTVs. Missense mutations should be located in hotspot locations previously reported in COSMIC in at least a few cancers. For instance, *FBXW7* is found recurrently mutated in this study, most often at known hotspot positions. However, a few missense mutations detected in this gene were reported in a low number of intestinal cancers. Even less missense mutations were not reported and could therefore not be considered genuine driver mutations. The frequency of driver mutations was compared between samples taken from of MutSα and MutLα deficient tissues. These frequencies were similarly compared to the driver mutation landscape of 445 MMR-proficient healthy intestinal crypts (Lee-Six et al., 2019), which were annotated and filtered using the same approaches.

### Genomic Distribution Mutations

Genomic regions were masked based on the ENCODE blacklist to rule out errors due to read misalignment, regions of poor alignment or regions exhibiting anomalous coverage intensity signals in an approach similar as previously published (Supek and Lehner, 2015). The remainder of the unmasked genome was segregated into non-overlapping 100 kb (Figure S4B) or 1 Mb (Figures 6A-B and Figure S4C) windows and those for which a significant proportion (*≥* 50%) fails the above criteria are removed from further analysis. The Y chromosome was similarly excluded from analysis. This masking process yielded 26396 100 kb or 2588 1 Mb windows passing the above criteria. Different from previous approaches (Supek and Lehner, 2015), exonic areas are included. SBS densities were calculated per donor and tissue combination. For instance the normal colonic epithelium of case PD40102. SBS can be aggregated across intestinal crypts under the assumption that mutation rates are approximately similar across the different intestinal crypts from the same tissue. SBS are extracted from the phylogenetic trees to prevent mutations being duplicated. For each donor and tissue combination, the SBS density of each window is determined by aggregating the SBS across all microdissected intestinal crypts and dividing the total count by the effective window length after masking. In the end there are 26396 or 2588 SBS density values for each donor and intestinal tissue combination. The relative mutation frequency (RMF) is determined by dividing the SBS density value by the mean SBS density value across all windows per donor and tissue combination.

A DNA replication timing profile was generated from 11 Repli-seq datasets from ENCODE (Consortium, 2012) downloaded through the UCSC genome browser (hg19). The data comprises genome-wide wavelet-smoothed Repli-seq values per 1kb bins for 11 cell lines (BG02ES, BJ, GM12878, Hela-S3, HepG2, HUVEC, IMR90, K562, MCF-7 and NHEK). The median value across all cell lines was calculated per 1kb window and from these median values the median value was calculated for 100 kb or 1 Mb non-overlapping windows. These non-overlapping windows were sorted based on the summarized Repli-seq value and distributed among 5 equally-sized domains, named the Earliest, Early, Median, Late and Latest replication timing domains. The higher the summarized Repli-seq value the earlier it is expected that this region of the genome undergoes replication. Thereafter mutation windows are assigned to the replication timing domains to assess the effect of replication timing on the relative SBS mutation rate.

For each non-overlapping RMF window the density of coding sequence is determined with the EnsDb.Hsapiens.v75 package in R, while the CpG density is determined using the BSgenome.Hsapiens.UCSC.hg19 package in R. RMF, replication timing, coding genome density and CpG density were plotted with Gviz (Hahne and Ivanek, 2016).

### Relative Mutation Rate

Whole genome bisulfite sequencing (WGBS) data from 2 normal sigmoid cell colon samples were downloaded from GEO Omnibus (GSM983645 and GSM1010989) and was used for assessing the methylation state of CpGs. The analysis was restricted to CpG sites covered by at least 1 WGBS sample, the average is calculated in case sufficient coverage was present in both samples. The relative mutation rate for CpG or 5mC can be calculated by determining the CpG abundance (RMR-CG) or 5mC (RMR-5mC) abundance as previously described (Sanders et al., 2018). RMR-CG reflects the number of mutations expected in 1Mb of CpGs in a particular genomic feature if 1000 somatic mutations are randomly selected from the mutation catalogue:

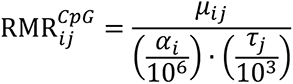

Where *μ*_*ij*_ represents the <u>C</u>G><u>T</u>G mutation count in feature i for sample j, *α_i_* represents the total number of CpGs in feature i, and *τ_j_* is the total <u>C</u>G><u>T</u>G mutation burden for sample j.

For RMR-5mC the *α_i_* is replaced with the methylation level sum across feature i:

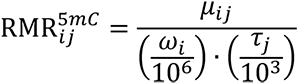

Where:

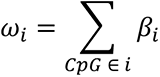

Where *β_1_* is the average methylation level (ranging between 0 – no methylation – and 1 – full methylation). RMR-5mC corrects for the methylation state of features of interest. For instance, promoters often have little methylation.

To assess the linear relationship between methylation level and mutation probability, <u>C</u>G><u>T</u>G mutations are binned per sample based on the methylation level detected in the normal sigmoid colon into consecutive bins of 0.1 interval, ranging from 0 (no methylation) to 1 (full methylation) with a 0.1 interval. The total count per bin is divided by 1.000.000 and the age of the donor to assess the annual mutation rate per million methylated CpGs.

### Fitting Mutational Signatures for Regions of Interest

For normal intestinal epithelium crypts mutations per region of interest are determined. First, replication timing domains (Early, Earliest, Median, Late, Latest) as described before. Second, across the coding sequence (CDS), all human exons, complete genes and the intergenic space. Mutational signatures extracted from the microdissected samples, described under ‘Mutational signature extraction’, are fitted to the mutational catalogues of the replication domains or regions of interest with TensorSignatures (Vöhringer et al., 2020). For MutSα-deficient intestinal crypts the contribution of MMR-1 is high in early replication regions and the CDS due to the increase CpG density in these regions. A beta generalized linear mixed model was fitted to test whether MutSα-deficient intestinal crypts have an enrichment of MMR-1 in these regions compared to the MutLα-deficient intestinal crypts. Fitting was done with glmmTMB (Brooks et al., 2017) using the exposure of MMR-1 (proportion) as the dependent variable, MutSα-deficiency as fixed effect (dichotomous variable) and donor as random intercept (multiple samples per tissue). The 95%-CI was determined by 10.000 bootstraps via bootMer from lme4, while the p-value was determined by parametric bootstrapping for 10.000 bootstraps with a full model, containing MutSα-deficiency as fixed effect and a reduced model, without MutSα-deficiency as variable and calculating the loglikelihood deviance as comparator.

### Assessing MMRD Genotype

MSI status for endometrial, colorectal and stomach cancer from The Cancer Genome Atlas (TCGA) was downloaded from the GDC data portal. For the remainder of the TCGA cohort, including endometrial, colorectal and stomach cancers with a negative MSI status, samples were selected with a high whole exome sequencing (WES) mutation burden (*≥* 150 somatic mutations) for further characterization. The mutational catalogues of selected WES were fitted with the standard COSMIC v3 MMR signatures SBS-6, SBS-15, SBS-20, SBS-21 and SBS-26 and SBS-1. Cases with an exposure of *≥* 0.25 for one MMR signature or SBS-1 were selected for genotyping. Mutational profiles were on occasion manually reviewed to ensure correct cases were selected for further analysis. Including the constitutive mismatch repair deficiency (CMMRD) this resulted in 337 possible MMRD cancers.

Tumor-only variant calling was done with Mutect2 (Benjamin et al., 2019). Protein truncating variants (PTV) in *MLH1*, *MSH2*, *MSH6* or *PMS2* were considered disruptive. Missense mutations in these genes in a hypermutation setting can result in incorrect genotyping. Only missense mutations listed as ‘likely pathogenic’ or ‘pathogenic’ in the InSiGHT (Thompson et al., 2014) or ClinVar (Landrum et al., 2018) databases were considered clinically relevant. The remainder of missense mutations in the MMR genes were only considered relevant if there was strong evidence in the literature from *in vivo*/*in vitro* assays that a hypermutation phenotype is observed. Transcript fusions or structural variants effecting a MMR gene was considered relevant. Finally, CNV loss as determined by ASCAT (Van Loo et al., 2010) or by local loss of coverage was considered relevant.

Based on sample purity as estimated by ASCAT, cases were only considered genotyped, if the following is found: a clinically relevant homozygous mutation after taking purity into account, compound heterozygous relevant mutations in the same gene, digenic relevant mutations or when a clinically relevant mutation is combined with a CNV or SV. Promoter methylation followed by gene silencing, commonly observed for *MLH1* in CpG island hypermethylation (CIMP) colorectal cancers, was not taken along because the methylator phenotype might be an event secondary to neoplastic change or is transient in nature. This resulted in 117 MMRD cancers being genotyped. For a considerable number of cases the clinically relevant variant is germline, presumably Lynch syndrome. In accordance with TCGA procedures the cases have been anonymized in all figures and tables. The names reflect the TCGA cancer subtype only and additional information on the MMR genes with disruptive variants are included in the supplementary tables.

### Dirichlet Regression

The extracted mutational signatures are fitted to the TCGA and CMMRD WES mutational catalogues to determine the mutational signature composition per case. Dirichlet regression (Maier, 2014) was applied to relate changes in the mutational signature composition to the MMRD genotype or the deficiency of a specific MMR complex. Cases with clinically relevant variants in both the MutSα and the MutLα (digenic) complexes were excluded from the analysis. It remains difficult to assess whether the loss of both complexes in cancer has a compounding effect on a mutational level or whether there was an chronological order in the loss of both complexes. Cases with digenic clinically relevant mutations in the same complex, e.g., *MSH2* and *MSH6*, were retained. First, PMS2-deficient cancers were used as the reference when linking changes in the mutational signature composition to the MMRD genotypes. This analysis was performed on all the MMRD cancers fulfilling the listed criteria. Cancers with MutLα-deficiency were used as reference when comparing MutSα-deficiency to MutLα-deficiency for changes in the mutational signature composition. This analyses was performed across all MMRD cancers, all MMRD cancers commonly associated with MMRD (endometrial, colorectal, stomach and prostate cancer), endometrial cancer alone, colorectal cancer alone, stomach cancer alone and all other remaining MMRD cancers.

## SUPPLEMENTARY FIGURES

**Figure S1.**
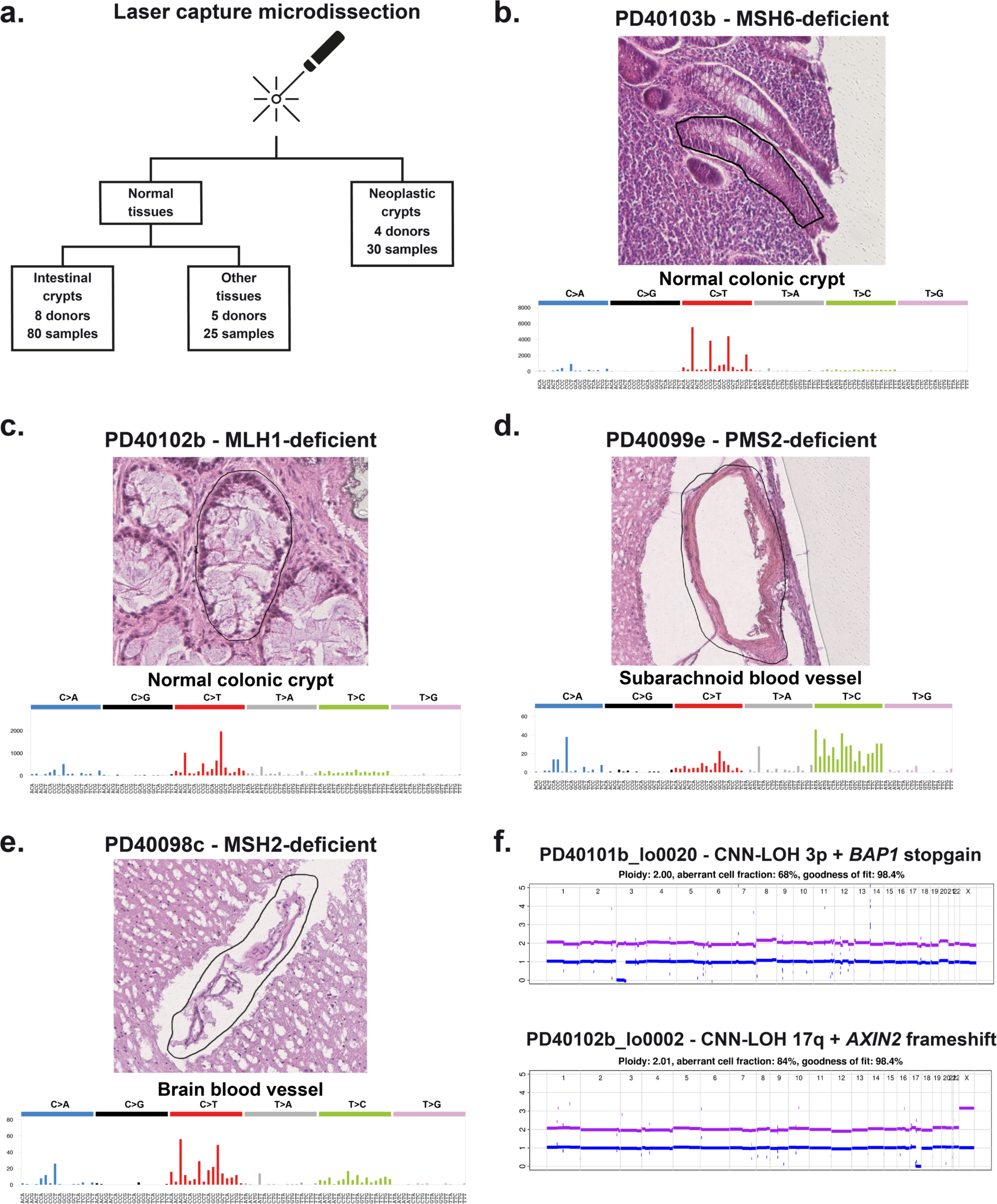
Mutational Profiles and Genetic Lesions Determined from CMMRD Tissues by Laser Capture Microdissection, Related to Figures 1-2 and 5. (**a**) Sample collection outline for CMMRD tissues by laser capture microdissection. (**b-e**) Microdissected areas and matched mutational profile for (**b**) a normal colonic crypt from MSH6-deficient donor PD40103, (**c**) a normal colonic crypt from MLH1-deficient donor PD40102, (**d**) a subarachnoid blood vessel collected from a brain sample of PMS2-deficient donor PD40099 and (**e**) a blood vessel collected from a brain sample from MSH2-deficient donor PD40098. (**f**) Examples of homozygous loss of driver genes through truncating mutations coinciding with copy number neutral loss of heterozygosity in individual crypts of donors PD40101 and PD40102.

**Figure S2.**
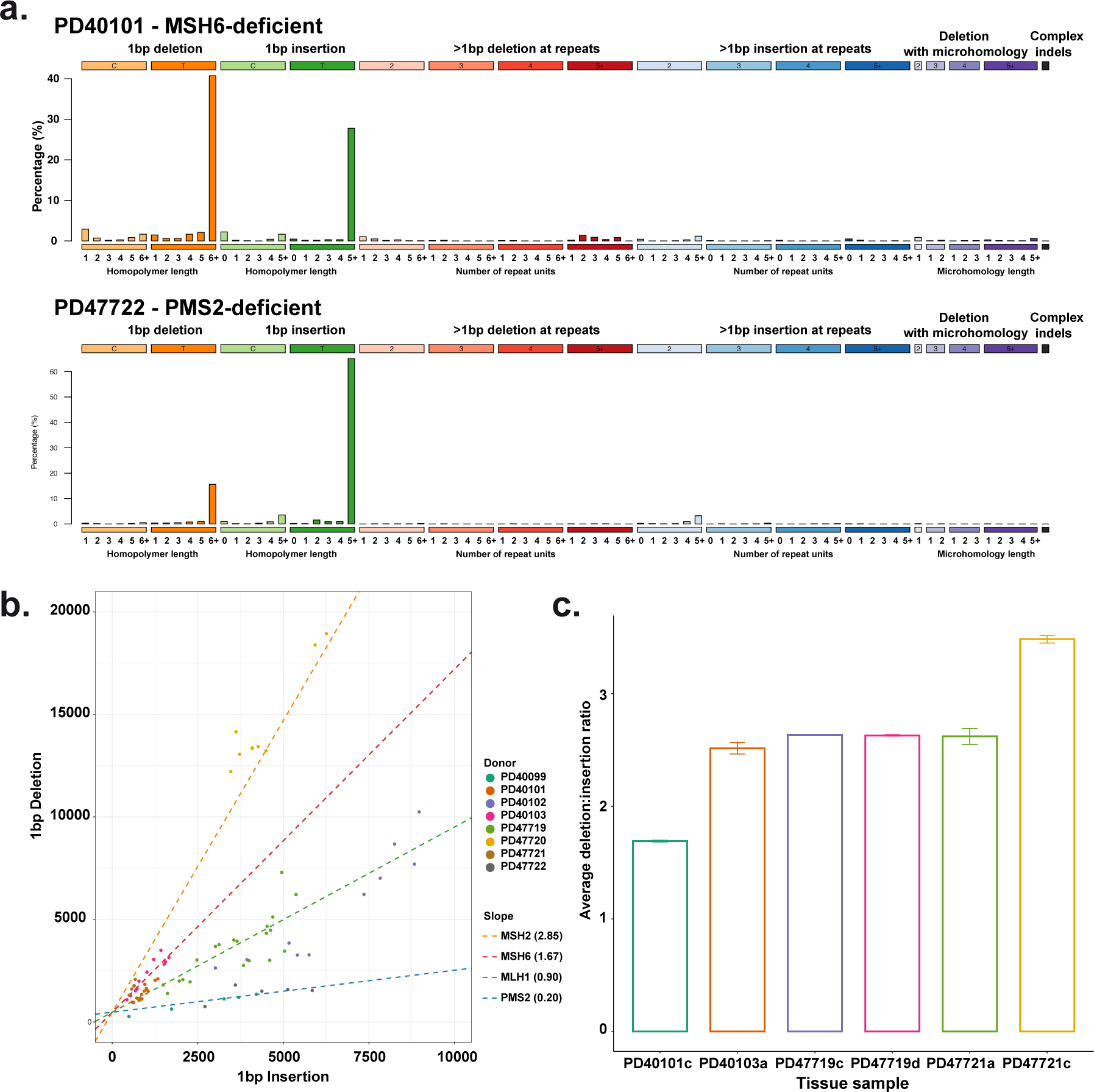
Insertion and Deletion Types and Acquisition Rates Differs Across MMRD Genotype, Related to Figures 1 and 3-4. (**a**) Typical insertion and deletion (indel) classification profiles observed from normal intestinal crypts from a MSH6-deficient and PMS2-deficient donor. Similar indel classification profiles were observed for other donors with the same MMRD genotype. (**b**) Linear mixed model fit for 1bp insertions against 1bp deletions with the MMRD genotype (MSH2, MSH6, MLH1 and PMS) included as fixed effect, donor included as random intercept and MMRD genotype included as random slope. (**c**) Average 1bp deletion-insertion ratios calculated across intestinal crypts from collected neoplastic tissues.

**Figure S3.**
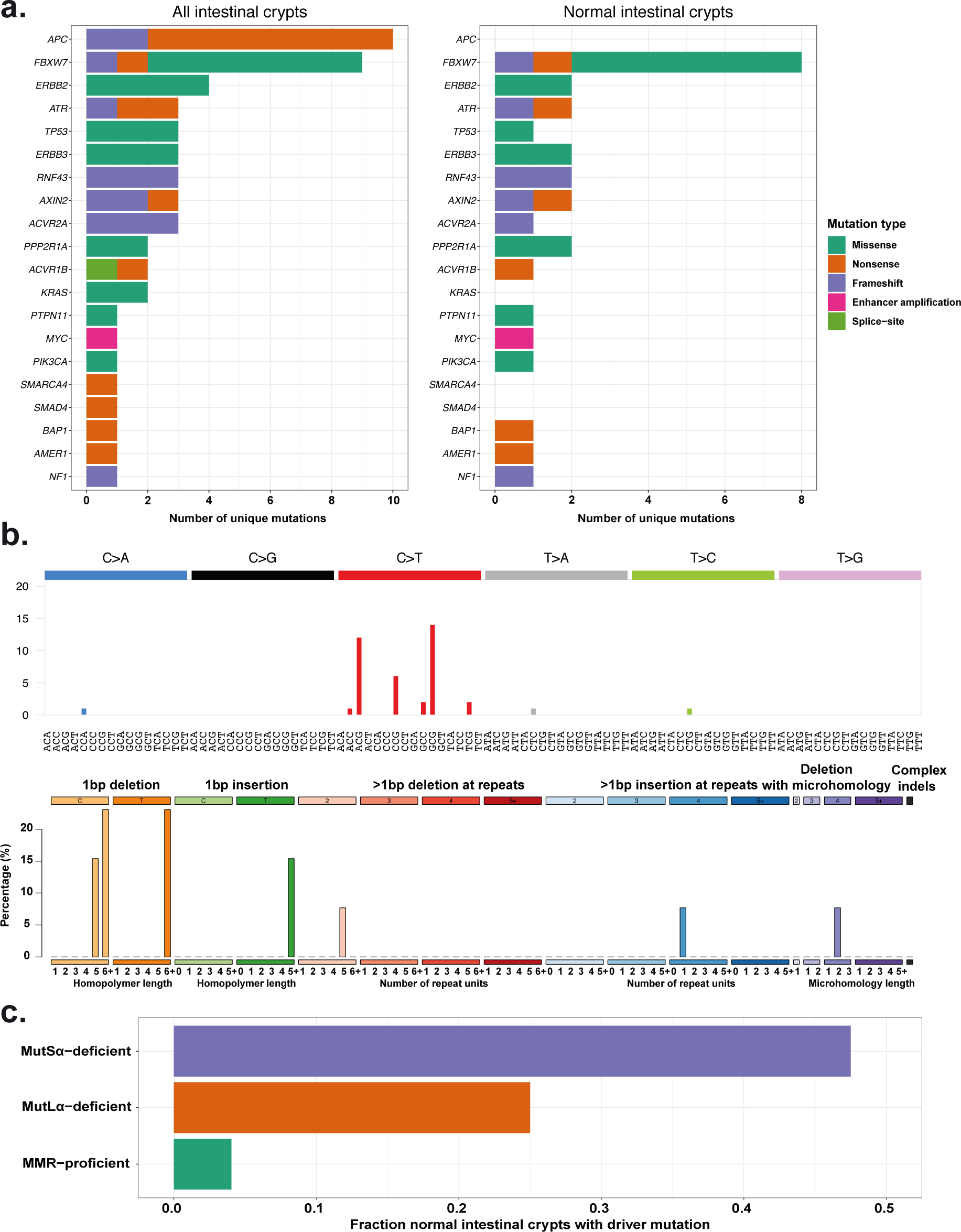
MMRD Driver Mutation Landscape Recapitulates Active Mutational Processes and Is Enriched Compared to the Healthy Population, Related to Figures 3-4. (**a**) Driver mutation count coloured by mutation type for (Left) all intestinal crypts isolated and (Right) all normal intestinal crypts. Mutation types include: missense, nonsense, essential splice site, frameshifts mutations and copy number changes. (**b**) SBS and indel mutational profiles of driver mutations detected in healthy and neoplastic intestinal crypts. (**c**) Fraction of normal intestinal crypts with driver mutations as detected in MutSα-deficient donors (MSH2 or MSH6, n = 40 crypts), MutLα-deficient donors (MLH1 or PMS2, n = 40 crypts) and MMR-proficient donors (n = 445 crypts).

**Figure S4.**
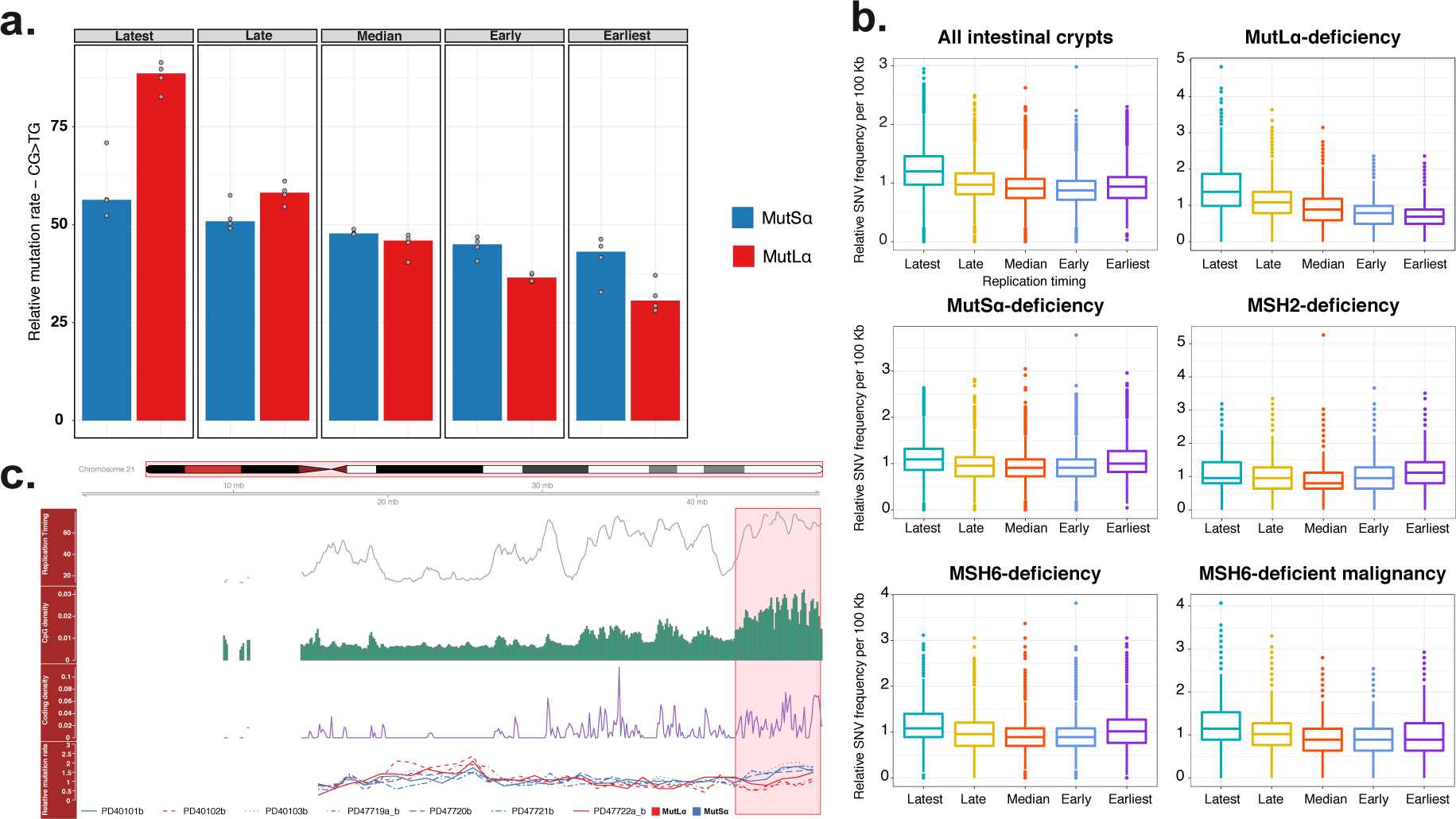
Genomic Distribution of Mutations in Intestinal Crypts is Associated with MMRD Genotype and Disease State, Related to Figures1, 3-4 and 6. (**a**) Relative mutations rates (RMR) of <u>C</u>G><u>T</u>G mutations was calculated for the replication timing domains (latest, late, median, early and earliest) by correcting for CpG-density and sample-specific mutation burden. Calculated RMR values were grouped based on MutSα or MutLα-deficiency to illustrate the changes in <u>C</u>G><u>T</u>G mutation rates across replication timing domains. (**b**) Relative SBS frequency boxplots for 100 kb non-overlapping bins across the genome grouped on replication timing domains. Data for healthy intestinal crypts is illustrated for all donors, donors grouped on MMR complex deficiency and loss of individual MMR components. Data for MSH6-deficient neoplastic intestine is illustrated in the bottom-right panel. (**c**) Distribution of mutations and established determinants of mutation rate for chromosome 21 across non-overlapping 1 Mb bins. From top to bottom. (I) Average replication timing profile. High values indicate early replication and vice versa. (II) CpG-dinucleotide density. (III) Coding genome density. (IV) Average relative SBS frequency of healthy intestinal crypts for individual donors as indicated by line type. Line color indicates the MMR complex lost for each donor. Highlighted areas indicate broad regions of high CpG and coding genome density.

**Figure S5.**
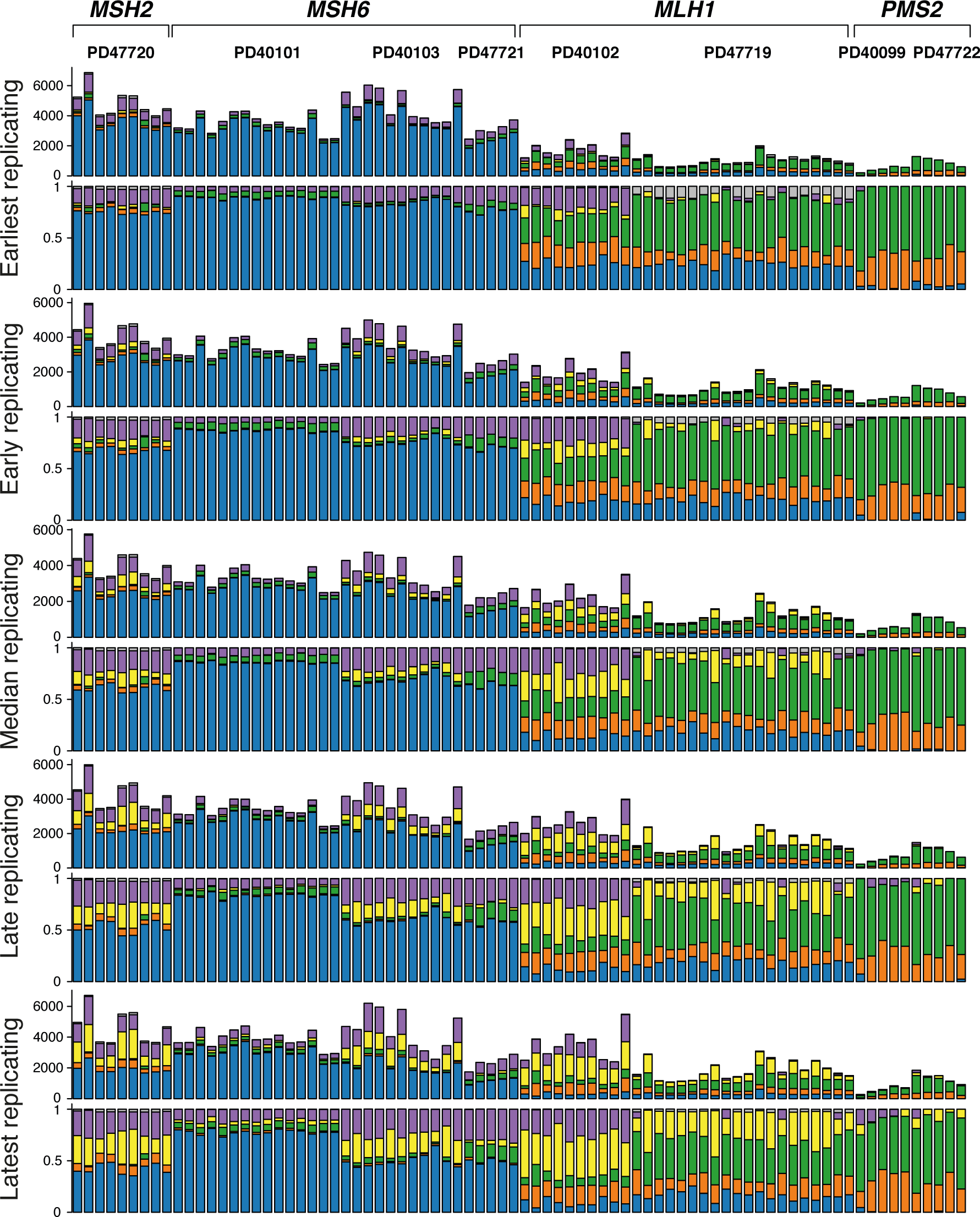
Differential DNA Damage Acquisition Across Replication Timing Domains Reveals the DNA Repair Patterns of Individual MMR components, Related to Figures 2-4 and 6. Mutations where counted across the replication timing domains. Mutational signatures extracted from the genome-wide normal CMMRD intestinal crypt mutation catalogues (Figure 2) were fitted to each replication timing domain to establish mutational signature composition per sample and domain. Samples are grouped by donor and MMR complex deficiency. From top to bottom: Mutational signature attribution based on all mutations (SBSs and indels) and the relative mutational signature attribution for each replication timing domain, ranging from earliest, early, median, late, latest.

**Figure S6.**
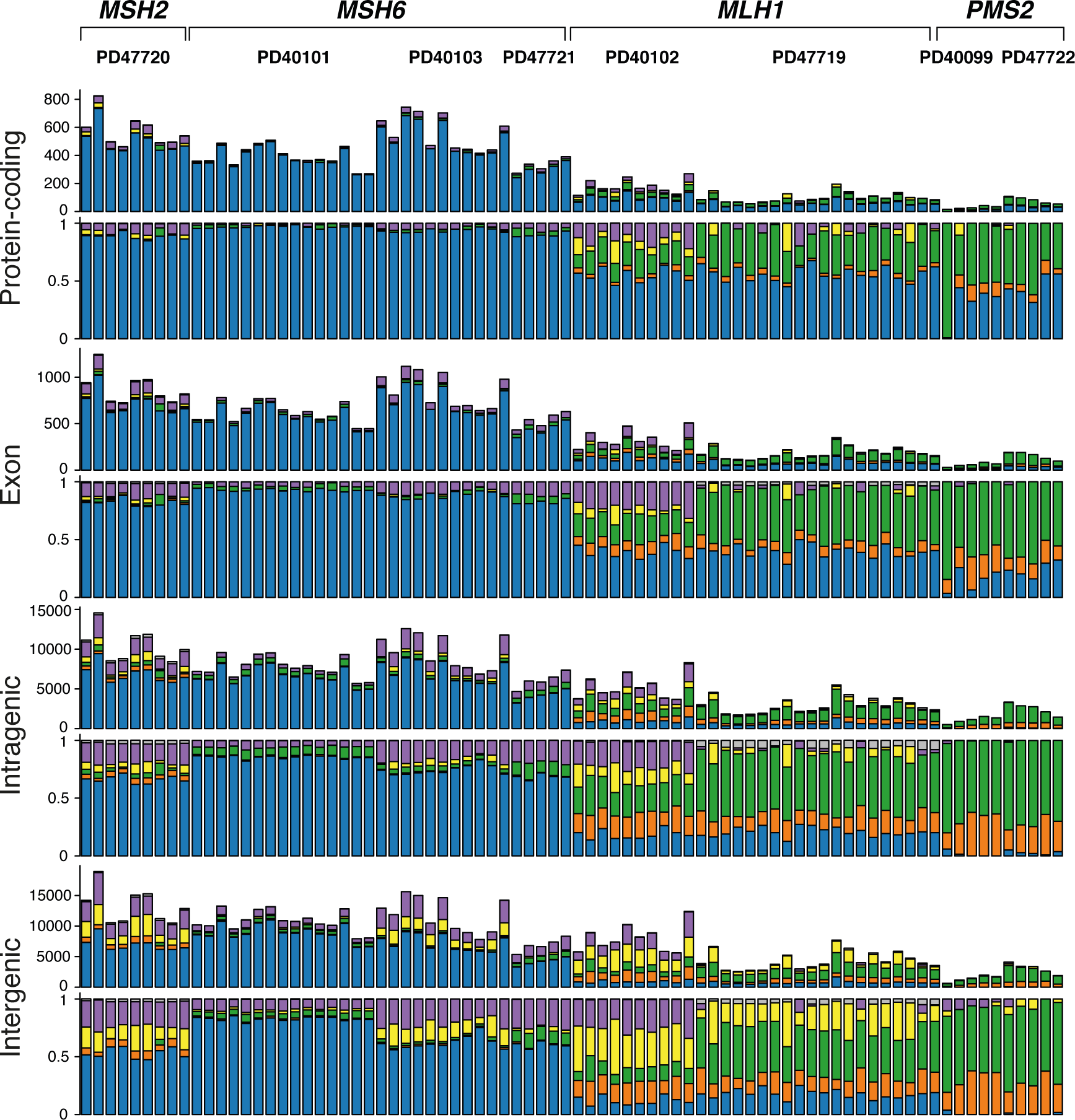
Differential DNA Damage Acquisition Across Genomic Elements Reveals the DNA Repair Patterns of Individual MMR components, Related to Figures 2-4 and 6. Mutations were counted across different genomic elements, ranging from the coding sequence, all exons (including untranslated regions), all genic constituents (exons and introns) and the intergenic space. This order of genomic elements follows the trend of methylated CpG density. Mutational signatures extracted from genome-wide mutation catalogues (Figure 2) were fitted to each genomic element to establish the mutational signature composition per sample and genomic element. Samples are grouped by donor and MMR complex deficiency. From top to bottom: Mutational signature attribution based on all mutations (SBSs and indels) and the relative mutational signature attribution for each genomic segment.

**Figure S7.**
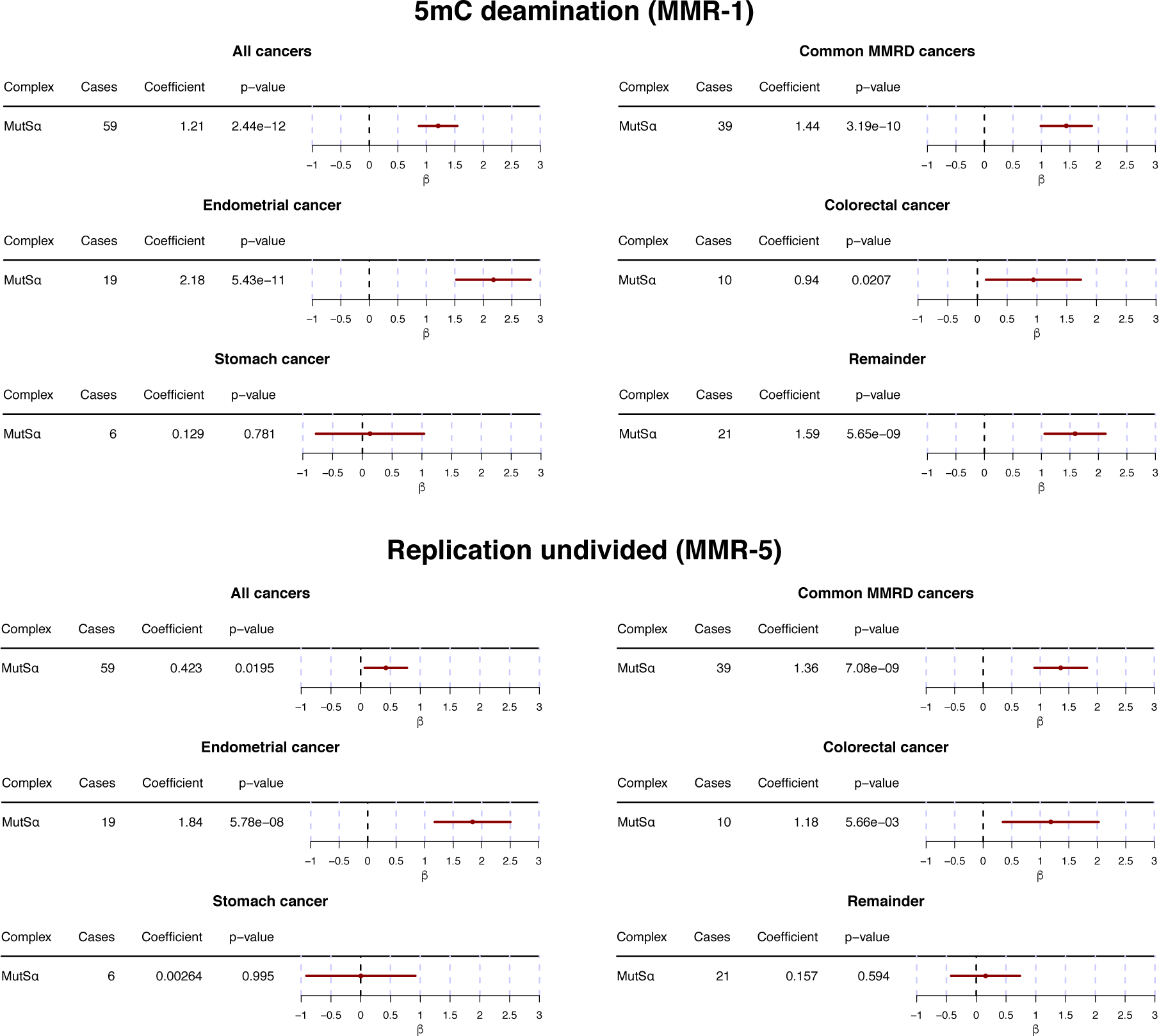
Validation of Mutational Signature Activity in Public Cancer Datasets, Related to Figures 2 and 7. Dirichlet regression performed on the mutational signature composition established by fitting the extracted mutational signatures to the WES mutational catalogues of MMRD cancers. Cases with loss of both MMR complexes (MutSα and MutLα) were excluded from the analysis. Cancers with MutLα-deficiency were used as reference. Enrichment or significance should be considered relative. Significance of mutational signature enrichment was tested in all MMRD cancers, the cancer subtypes enriched for MMRD (endometrial, colorectal, stomach and prostate cancer), endometrial cancer, colorectal cancer, stomach cancer and the remainder of MMRD cancers. (**a**) Validation of the association between 5mC deamination signature (MMR-1) activity and MutSα-deficiency in all predefined cancer categories with the exception of MMRD stomach cancer. (**b**) Validation of the association between ‘Replication Undivided’ signature (MMR-5) activity and MutSα-deficiency in all predefined cancer categories with the exception of MMRD stomach cancer and the remainder of MMRD cancers.

